# Structure of *C. elegans* TMC-1 complex illuminates auditory mechanosensory transduction

**DOI:** 10.1101/2022.05.06.490478

**Authors:** Hanbin Jeong, Sarah Clark, April Goehring, Sepehr Dehghani-Ghahnaviyeh, Ali Rasouli, Emad Tajkhorshid, Eric Gouaux

**Affiliations:** Vollum Institute, Oregon Health & Science University, Portland, Oregon 97239, USA; Howard Hughes Medical Institute, Oregon Health & Science University, Portland, Oregon 97239, USA; Theoretical and Computational Biophysics Group, NIH Center for Macromolecular Modeling and Bioinformatics, Beckman Institute for Advanced Science and Technology, Department of Biochemistry, and Center for Biophysics and Quantitative Biology, University of Illinois at Urbana-Champaign, Urbana, Illinois 61801, USA

## Abstract

The initial step in the sensory transduction pathway underpinning hearing and balance in mammals involves the conversion of force into the gating of a mechanosensory transduction (MT) channel. Despite the profound socioeconomic impacts of hearing disorders and the fundamental biological significance of understanding MT, the composition, structure and mechanism of the MT complex has remained elusive. Here we report the single particle cryo-EM structure of the native MT TMC-1 complex isolated from *C. elegans*. The 2-fold symmetric complex is composed of 2 copies each of the pore-forming TMC-1 subunit, the calcium-binding protein CALM-1 and the transmembrane inner ear protein TMIE. CALM-1 makes extensive contacts with the cytoplasmic face of the TMC-1 subunits while the single-pass TMIE subunits reside on the periphery of the complex, poised like the handles of an accordion. A subset of particles in addition harbors a single arrestin-like protein, ARRD-6, bound to a CALM-1 domain. Single-particle reconstructions and molecular dynamics simulations show how the MT complex deforms the membrane bilayer and suggest crucial roles for lipid-protein interactions in the mechanism by which mechanical force is transduced to ion channel gating.

## Main

The auditory system is endowed with a remarkable ability to detect a wide range of acoustic wave frequencies and amplitudes by transducing vibrational mechanical energy into membrane potential depolarization, followed by signal processing in higher brain centers, thus enabling the sensation of sound^1^. Dysfunction of the auditory system, from injury, environmental insult or genetic mutation, is associated with age-related hearing loss. Hearing impairment and deafness impacts over 460 million individuals worldwide, with an estimated annual cost of unaddressed hearing loss at $750-790 billion ^2^. Input into the auditory and the closely related vestibular system, like other sensory systems, is initiated by receptor activation on peripheral neurons. Despite intense investigation over several decades, the molecular composition, structure and mechanism of the receptor for mechanosensory transduction, deemed the MT complex, has remained unresolved.

Multiple lines of investigation from studies on humans and model organisms including mice, zebrafish and *C. elegans*, nevertheless, have shed light on the proteins that form the MT complex and their likely roles in its function ^3^. These include the tip link proteins, protocadherin-15 and cadherin-23, that in hair cells transduce the force derived from stereocilia displacement to the opening of the ion channel component of the MT complex ^4, 5^. The transmembrane ion channel like proteins 1 and 2 (TMC-1 and TMC-2 are the likely pore-forming subunits of the MT complex, candidates that first came to prominence from human genetic studies ^6^, and more recently gained traction as the ion conduction pathway via biophysical and biochemical investigations ^7, 8^. Additional proteins, some of which may be ‘auxiliary subunits’, have been associated with either the biogenesis or function of the MT complex and include transmembrane inner ear protein (TMIE) ^9–11^, Ca^2+^ and integrin binding protein 2 (CIB2) ^12–14^, lipoma HMGIC fusion-like protein 5 (LHFPL5) ^15–17^, transmembrane O-methyl transferase (TOMT) ^18, 19^ and possibly ankyrin ^13^.

Isolation of the MT complex from vertebrate sources or production of a functional complex via recombinant methods have proven unsuccessful. Complex purification from native sources is particularly challenging due to the small number of complexes per animal, estimated as ∼3 × 10^6^ per mammalian cochlea ^20^, miniscule compared to the number of photoreceptors of the visual system, which is ∼4 × 10^14^ per murine eye ^21^. To surmount challenges with vertebrate MT complex availability, we turned to *C. elegans*, an animal that harbors a MT complex used for sensing tactile stimuli. We note first that *C. elegans* expresses crucial components of the vertebrate MT complex, including the TMC-1 and TMC-2 proteins, in addition to a CIB2 homolog, known as CALM-1, as well as TMIE ^13^. Second, worms that are devoid of TMC-1 exhibit attenuated light touch responses ^13^. Third, despite the limited expression of the TMC proteins in *C. elegans*, it is feasible to grow a sufficient number of worms to isolate enough complex for structural studies. We thus modified the *C. elegans tmc-1* locus by including a fluorescent reporter and an affinity tag, thereby allowing us to monitor expression via whole animal fluorescence and fluorescence-detection size-exclusion chromatography (FSEC) ^22^, and to isolate the TMC-1 complex by affinity chromatography. Together with computational studies, we elucidated the composition, architecture and membrane interactions of the complex, and suggest mechanisms for gating of the ion channel pore by both direct protein interactions and via the membrane bilayer.

### TMC-1 complex is a dimer

We generated a transgenic knock-in worm line where a nucleic acid sequence encoding an mVenus- 3xFLAG tag was inserted at the 3’ end of the TMC-1 coding sequence, immediately before the stop codon (Supplementary Fig. 1). The engineered, homozygous worm line, deemed *tmc-1::mVenus*, was characterized by spectral confocal imaging, revealing mVenus fluorescence in the head and tail neurons, and in body wall and vulval muscles (Fig. 1a), consistent with previous studies demonstrating expression of TMC-1 in these cells ^23^. The TMC-1 complex was isolated from the *tmc-1::mVenus* transgenic worms by affinity chromatography and further purified by size exclusion chromatography (SEC) (Fig. 1b). The estimated molecular weight of the TMC-1 complex by SEC is ∼780 kDa, suggesting that the complex harbors multiple TMC-1 protomers and perhaps additional, auxiliary subunits.

**Fig. 1.**
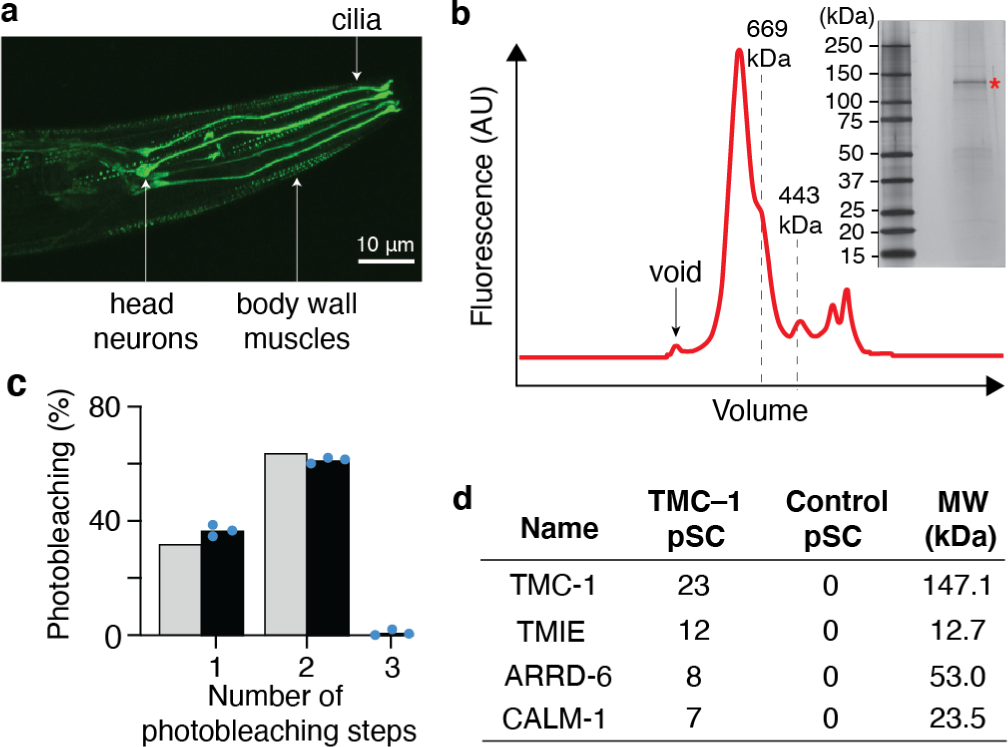
Dimeric TMC-1 complex from *C. elegans* copurifies with additional proteins. **a,** Spectral confocal image of mVenus fluorescence in an adult *tmc-1::mVenus* worm showing mVenus fluorescence in the head neurons, cilia and body wall muscles. Shown is one representative image of five total images. **b**, Representative FSEC profile of the TMC-1 complex, detected via the mVenus tag. Inset shows a silver-stained, SDS-PAGE gel of the purified TMC-1 complex. Red asterisk indicates TMC-1. **c**, The distribution of mVenus photobleaching steps for the TMC-1 complex is consistent with a binomial function (grey bars) an assembly with two fluorophores. A total of n = 600 spots were analyzed from three photobleaching movies (200 spots per movie). Each movie is represented by a blue dot. **d**, Analysis of TMC-1 complex by mass spectrometry (MS) shows selected identified proteins in order of decreasing peptide spectral counts. Proteins that were identified in the TMC-1 sample, but not in the control sample from wild-type worms, are shown. A full table can be found in Extended Data Fig. 2.

To independently interrogate the oligomeric state of the complex, we performed single molecule pulldown (SiMPull) experiments ^24^. Photobleaching traces of captured TMC-1 complexes demonstrate that ∼62% of the mVenus fluorophores bleached in two steps, 37% bleached in one step, and 1% bleached in three steps (Fig. 1c; Extended Data Fig. 1), consistent with the conclusion that within the TMC-1 complex there are two copies of the TMC-1 subunit. The discrepancy between the predicted ∼300 kDa molecular weight of a *C. elegans* TMC-1 dimer and the molecular weight of the complex estimated by SEC points towards the presence of auxiliary proteins. As several TMC-1 binding partners have been identified in worms ^13^ and in vertebrates ^3^, we next probed the composition of the TMC-1 complex using mass spectrometry (MS).

MS analysis of the TMC-1 complex identified three proteins that co-purified with TMC-1: (1) CALM-1, an ortholog of mammalian Ca^2+^ and integrin-binding family member 2 (CIB2); (2) an ortholog of mammalian transmembrane inner ear expressed protein (TMIE); (3) ARRestin domain protein (ARRD- 6), an ortholog of the mammalian arrestin-domain-containing family of proteins (Fig. 1d; Extended Data Fig. 2). All three proteins were found in the TMC-1 sample purified from transgenic worms but not in the control sample prepared from wild-type worms, consistent with their specific association with the TMC-1 complex. The mammalian ortholog of CALM-1 is CIB2, and CIB2 which together with TMIE are likely components of the mammalian MT complex, localize to stereocilia ^9–12, 14, 25^ and bind to heterologously expressed TMC-1 fragments through pull-down assays ^10^. By contrast, ARRD-6 has not been described as a component of either the *C. elegans* or vertebrate TMC-1 complexes. Despite repeated efforts, we found no evidence for the presence of UNC-44, the worm ortholog of mammalian ankyrin, in contrast with a previous report that UNC-44 forms a complex with CALM-1, is necessary for TMC-1 mediated mechanotransduction, and is the ‘gating spring’ of the TMC-1 complex ^13^, thus raising the question of the role of UNC-44 and by extension ankyrin to the structure and function of MT complexes in worms and vertebrates, respectively.

**Fig. 2.**
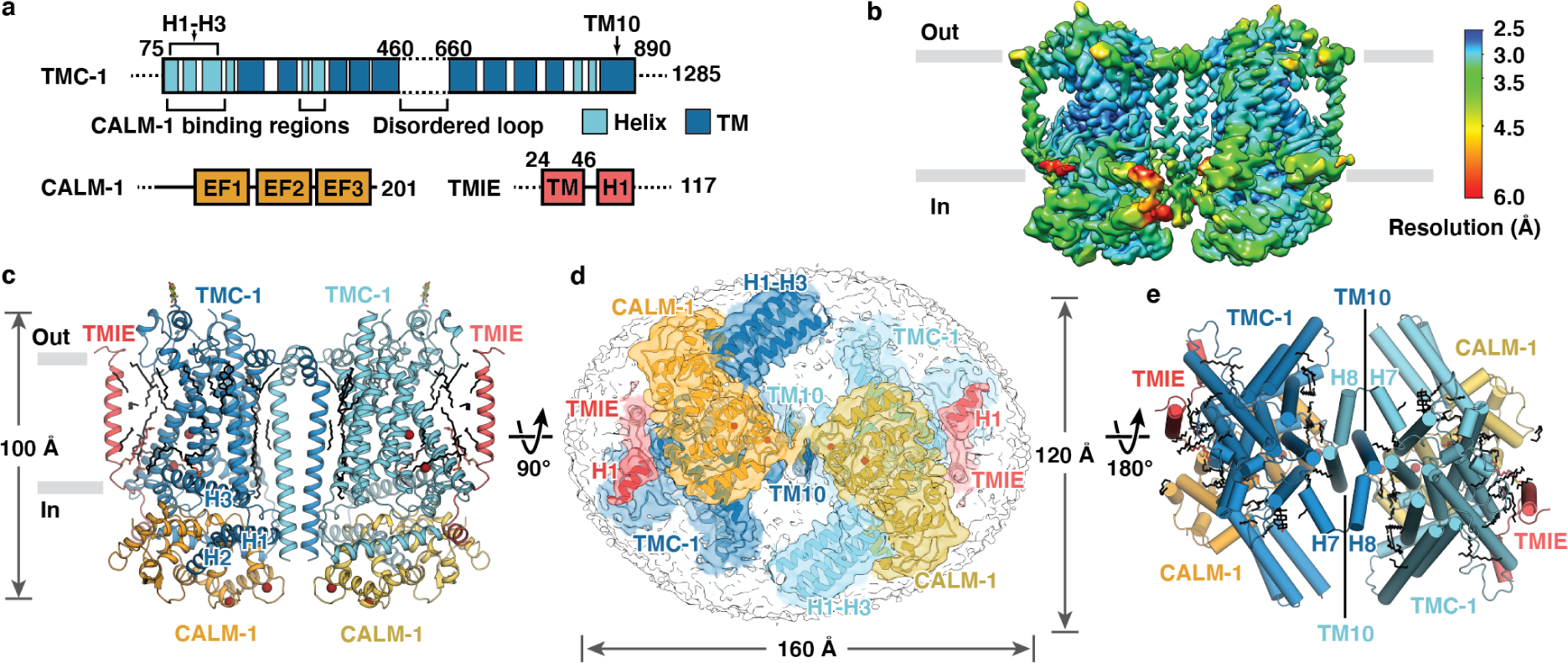
Architecture and subunit arrangement of the TMC-1 complex. **a**, Schematic representation of protein constructs that were isolated with TMC-1. **b,** Local resolution map of native TMC-1 complex after three-dimensional reconstruction. **c,** Overall architecture of native TMC-1 complex, viewed parallel to the membrane. TMC-1 (dark blue and light blue), CALM-1 (orange and yellow) and TMIE (red and pink) are shown in a cartoon diagram. Lipid-like molecules, *N*-Glycans, and putative ions are colored black, green, and dark red, respectively. **d,** Cytosolic view of the reconstructed map, fitted with the model. Subunit densities are colored as same in c) and the detergent micelle is shown in grey. **e,** A top-down extracellular view of the TMC-1 complex shows the domain-swapped dimeric interface. α- helices are represented as cylinders.

### Overall architecture of the TMC-1 complex

To elucidate the architecture and arrangement of subunits in the TMC-1 complex, we performed single particle cryo-electron microscopy (cryo-EM). TMC-1 is expressed at a low level in *C. elegans* and therefore approximately 6 × 10^7^ transgenic worms were required to yield ∼50 ng of TMC-1 complex for cryo-EM analysis. The TMC-1 complex was visualized on 2 nm carbon-coated grids that were glow discharged in the presence of amylamine. Cryo-EM imaging revealed a near ideal particle distribution and we proceeded to collect a dataset comprised of 26,055 movies. Reference-free 3D classification reconstruction, together with refinements, resulted in three well-defined classes (Extended Data Figs. 3-5, Extended Data Table 1). Two of these classes represent the TMC-1 complex in different conformational states, deemed the ‘Expanded’ (E) and ‘Contracted’ (C) conformations, both of which exhibit an overall resolution of 3.1 Å (Extended Data Figs. 3 and 5). A third class includes the auxiliary subunit ARRD-6 and was resolved at 3.5 Å resolution (Extended Data Fig. 4 and 5, Extended Data Table 1). Because the ‘E’ conformation has a few more distinct density features than the ‘C’ conformation, we will focus on the ‘E’ conformation in our initial discussion of the overall structure.

**Fig. 3.**
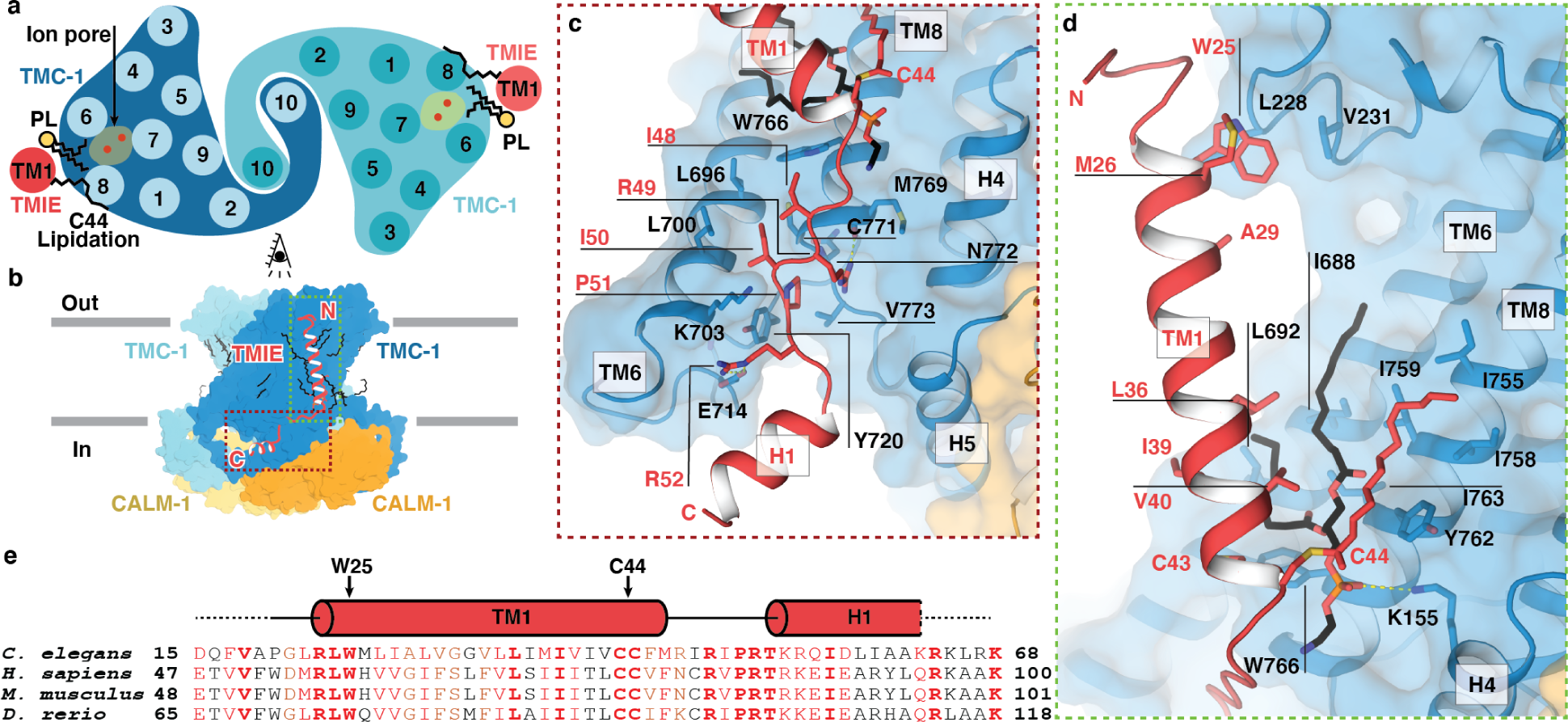
TMIE resides on the periphery of the TMC-1 complex. **a**, Schematic representation of TMC-1 (blue) and TMIE (red) transmembrane helices highlights the proximity of TMIE to the putative TMC-1 ion conduction pathway. Palmitoylation of TMIE C44 and phospholipids are shown in black. **b**, Overview of the interaction interface between TMIE and TMC-1, viewed from the side. **c**, Interface between the TMIE ‘elbow’ and TMC-1. Interacting residues are shown as sticks. **d**, Interface between TMIE transmembrane helix and TMC-1, highlighting key residues and lipids. Palmitoylation is shown in red and phospholipid is shown in black. **e**, Multiple sequence alignment of TMIE orthologs. Elements of secondary structure are shown above the sequences and key residues are indicated with black arrows.

**Fig. 4.**
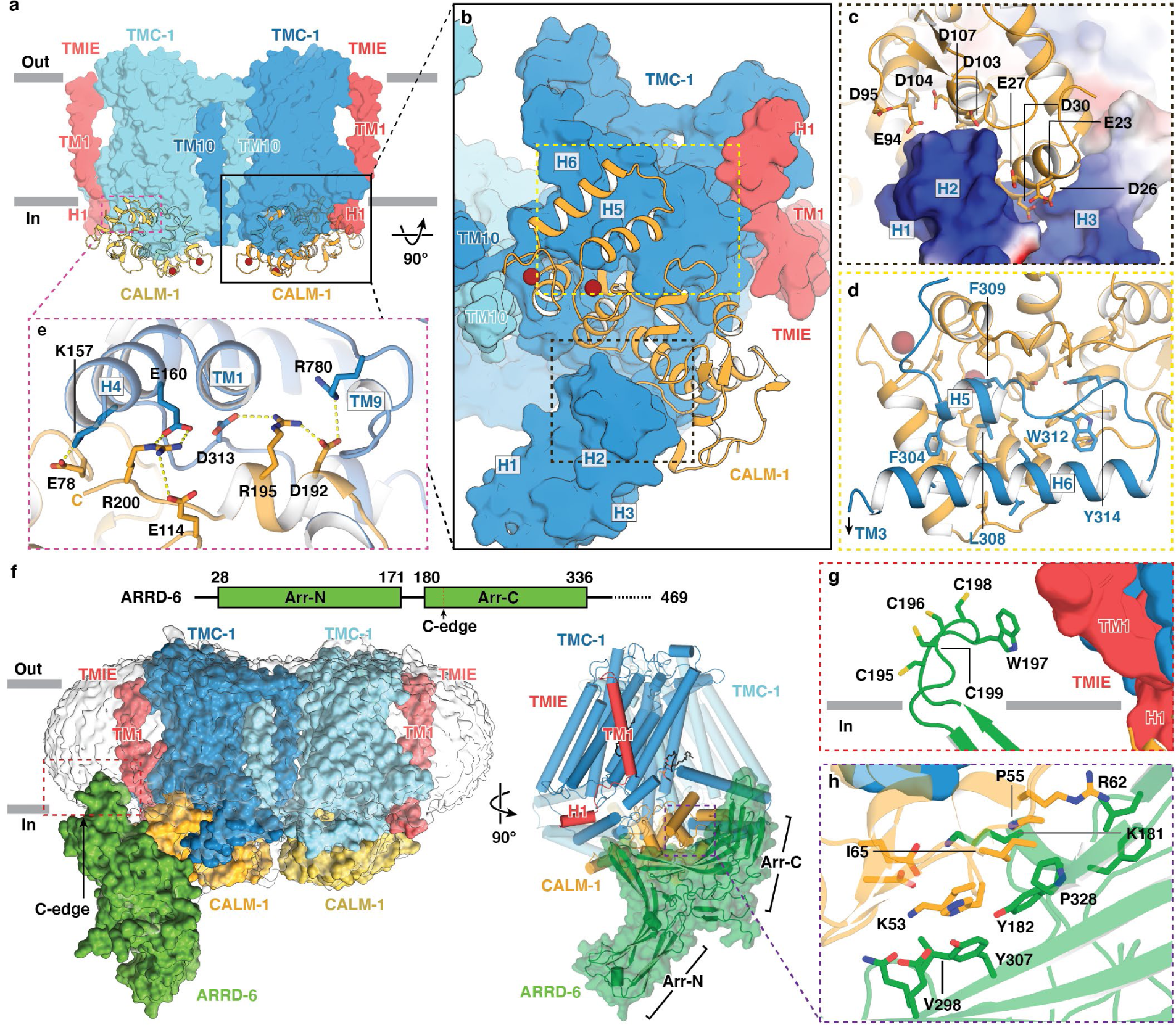
CALM-1 and ARRD-6 auxiliary subunits cap the cytoplasmic face of the TMC-1 complex. Binding interface between CALM-1 and TMC-1 viewed **a**, parallel to the membrane and **b**, perpendicular to the membrane. **c**, Binding interface between CALM-1 and TMC-1 H1-H3. The electrostatic surface of TMC-1 is shown, where blue represents basic regions and red represents acidic regions. CALM-1 is shown in yellow. **d**. Interface between CALM-1 and TMC-1 H5-H6. **e**, Salt bridges between the C-terminus of CALM-1 and TMC-1. Putative hydrogen bonds are shown as dashed lines. **f,** 3D reconstruction of the TMC-1 complex with ARRD-6 viewed parallel to the membrane. TMC-1, CALM-1, TMIE, and ARRD-6 are shown in blue, yellow, red, and green, respectively. A red dashed rectangle indicates the putative insertion site of the ARRD-6 C-edge loop into the micelle. A schematic diagram of ARRD-6 is shown above the reconstruction. **g**, Interface between the C-edge loop of ARRD-6 and the membrane. ARRD-6 residues that likely participate in membrane interactions are shown as sticks. **h**, Interface between ARRD- 6 (green) and CALM-1 (yellow), highlighting residues that are important for the binding interaction.

**Fig. 5.**
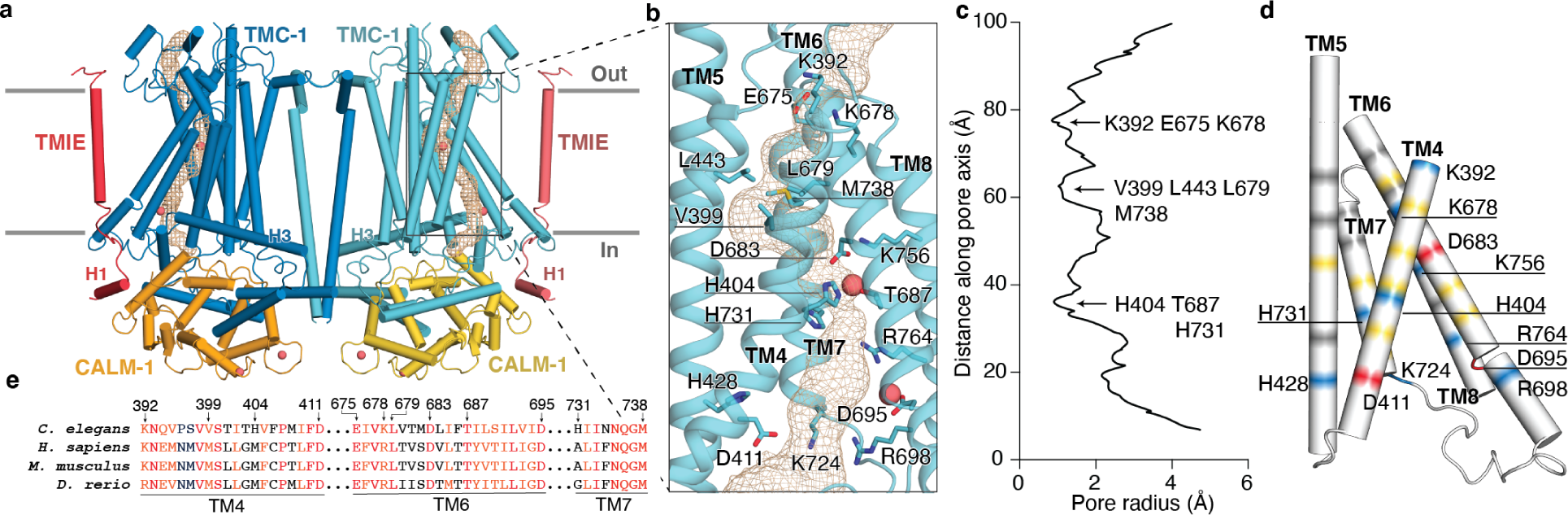
The putative ion conduction pore of TMC-1. **a**, The location of the pore (gold mesh) is shown in the context of the TMC-1 complex. Putative calcium ions are shown as pink spheres. **b**, An expanded view of the ion permeation pathway, highlighting pore- lining residues, shown as sticks, and putative ions (pink). **c**, van der Waals radius of the pore plotted against the distance along the pore axis, calculated by MOLE 2.0. **d**, Electrostatic potential of pore-lining residues are depicted in different colors: grey = nonpolar, yellow = polar, red = acid, blue = basic. Acidic and basic residues are labeled. **e,** Multiple sequence alignment of selected residues from TMC-1 TM4-TM7, four of the putative pore-forming helices.

The TMC-1 complex is dimeric in subunit stoichiometry, with a 2-fold axis of rotational symmetry centered at a site of contacts between the two TMC-1 subunits (Fig. 2). The transmembrane helices exhibit better local resolution than average, while disordered or dynamic peripheral components of the complex are resolved at lower resolution (Fig. 2b). When viewed perpendicular to the membrane plane, the complex has the shape of a ‘figure 8’, with TMC-1 subunits centered within the lobes of the ‘8’ (Fig. 2d). Each TMC-1 protomer consists of ten transmembrane helices with an overall arrangement that is reminiscent of the Ca^2+^-activated lipid scramblase ^26^, TMEM16 Cl^-^ channels ^27, 28^ and OSCA mechanosensitive ion channels ^29, 30^ (Extended Data Fig. 6). At the juncture of the figure ‘8’ lobes, the dimer interface is composed of domain-swapped TM10 helices (Fig. 2e), with contacts defined by van der Waals and electrostatic interactions, and by burial of 1,781 Å^2^ of solvent accessible surface area. Numerous well-ordered lipid molecules surround the transmembrane domain, many of which are intercalated in the grooves between transmembrane helices, and some of which are positioned at a large angle to the membrane plane.

**Fig. 6.**
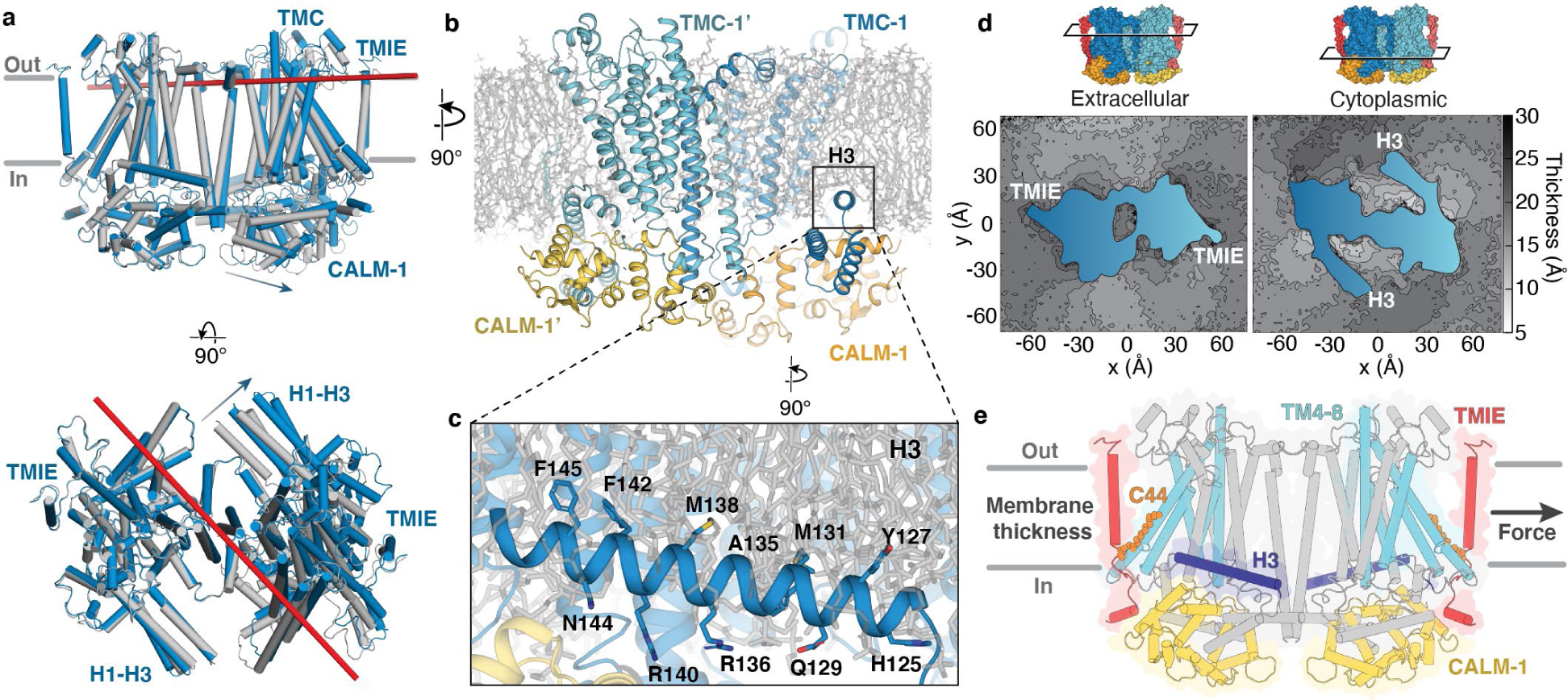
Conformational flexibility and membrane integration of the TMC-1 complex. **a**, A TMC-1 protomer from the ‘E’ conformation (blue) and the ‘C’ conformation (grey) were superposed based on backbone alpha-carbon atoms, revealing conformational differences in TMC-1, as well as CALM- 1 and TMIE. The axis of rotation is shown as a red bar and arrows indicate the direction of rotation from the ‘C’ to the ‘E’ state. **b**, MD simulation of the membrane-embedded TMC-1 complex shows deep penetration of the H3 helix into the lipid bilayer. **c**, Key residues that define the amphipathic nature of the H3 helix are shown as sticks. **d**, Lipid bilayer thickness for extracellular and cytosolic leaflets averaged over the last 500 ns of all three simulated replicas. The cross-section of the protein is shown in blue and the location of the cross section is indicated above the plots using a surface representation of the TMC-1 complex. Dark grey and light grey shades in the heatmaps represent membrane thinning and thickening, respectively. **e**, Schematic illustrating mechanisms by which direct or indirect forces are transduced to ion channel gating. TMIE is shown in red and palmitoylation of C44 is shown as orange spheres. The putative pore-forming helices of TMC-1 are shown in light blue, H3 is depicted in dark blue, and CALM-1 is yellow. Grey arrow (right) shows how membrane tension could directly gate the TMC-1 complex by exerting force on TMIE. Indirect force as a result of changes in membrane thickness could affect the position of the membrane embedded helix H3, modulating ion channel gating.

Poised to make extensive interactions with the inner leaflet of the membrane, the cytosolic domain harbors six helices oriented nearly parallel to the membrane. The two helices located closest to inner leaflet, H3 and H4, are amphipathic, a common feature among mechanosensitive ion channels ^31^ (Fig. 2c, d). The short linker between TMC-1 H3 and H4 is composed of nonpolar residues that interact with the inner leaflet membrane, forming hydrophobic contacts with the acyl chains of two lipids. The ∼400-residue, cytosolic C-terminus of TMC-1, which is predicted to be partially structured, was not visible in the cryo-EM map.

Because MS analysis of the purified MT complex identified nine peptides that spanned the entirety of the C-terminus, we suspect that this region is intact in the TMC-1 complex, but not visible due to conformational heterogeneity (Extended Data Fig. 2 legend).

Three partially structured loops decorate the extracellular side of the TMC-1 complex. Two of the loops are ∼60 residues in length, bridging TM1/TM2 and TM9/TM10, and are well-conserved between vertebrate and *C. elegans* TMC-1. Density features consistent with glycosylation can be found at N209, located in the loop between TM1 and TM2. By contrast, the ∼200-residue extracellular loop that connects TM5 and TM6 was not observed in the cryo-EM map. We detected two peptides from this region in the MS analysis (Extended Data Fig. 2 legend), indicating that the loop is present but not visible due to flexibility. This region is predicted to contain elements of secondary structure, as well as three predicted sites of *N-*linked glycosylation, but its function is unclear. The loop is not well conserved between TMC-1 and TMC-2, and its length in vertebrate TMC-1, at ∼50 residues, is substantially shorter.

Two additional subunits, CALM-1 and TMIE, present in two copies each, complete the ensemble of proteins associated with the ‘core’ TMC-1 complex. The quality of the cryo-EM map enabled unambiguous assignment of CALM-1 and TMIE auxiliary subunits (Fig. 2c) to density features of the TMC-1 complex map, in accord with the MS data. The CALM-1 subunits ‘grip’ the cytosolic faces of each TMC-1 protomer while each of the two TMIE subunits span the membrane, nearly ‘floating’ on the periphery of complex, flanking each TMC-1 subunit. CALM-1 makes extensive contacts with five of the six cytosolic helices, forming a ‘cap’ at the base of the TMC-1 transmembrane domain. By contrast, the TMIE subunits define the distal edges of the complex, participating in only a handful of protein-protein contacts on the extracellular and cytosolic boundaries of the membrane spanning regions, but with lipid mediated interactions through the transmembrane regions. Viewed parallel to the membrane and perpendicular to the long face of the complex, the arrangement of subunits resembles an accordion, with the TMIE transmembrane helices forming the instrument handles and the TMC-1 transmembrane domain defining the bellows (Fig. 2c).

### Extensive lipid-mediated interactions of TMIE with TMC-1

TMIE is an essential subunit of the vertebrate MT complex that is necessary for TMC-1 mediated MT in cochlear hair cells ^2^ and in zebrafish sensory hair cells ^10^. Multiple point mutations in TMIE are linked to deafness (Extended Data Fig. 7), and recent studies suggest a role for TMIE in TMC-1/2 localization and channel gating ^9–11, 32–34^. The *C. elegans* TMC-1 complex contains two copies of TMIE located on the ‘outside’ of each TMC-1 protomer (Fig. 3a). TMIE consists of a single transmembrane domain followed by an ‘elbow-like’ linker and a cytosolic helix (Fig. 3b). The flexible, positively charged C-terminal tail was not visible in the cryo-EM map. The interaction between TMIE and TMC-1 is mediated primarily by the cytosolic TMIE elbow, with the highly conserved R49 and R52 forming hydrogen bonds with backbone carbonyl atoms in TMC-1 TM6 and TM8, respectively (Fig. 3c, e). These arginine residues can be mapped to known deafness mutations in humans (R81C and R84W), highlighting the importance of these hydrogen bonds in the TMIE/TMC-1 interaction. Hydrophobic contacts between non-polar residues in the TMIE elbow and TMC-1 TM6 likely strengthen the complex. Additionally, W25 of TMIE, near the extracellular boundary, contacts L228 in the loop between TMC-1 TM1 and TM2 (Fig. 3d). Mutation of the corresponding residue in humans (W57) to a stop codon is a cause of deafness ^35^. We did not observe density for the N-terminal 17 residues of TMIE in the cryo-EM map and peptides from this region were not detected in the MS analysis, suggesting that the N-terminus contains a cleaved signal peptide (Supplementary Fig. 2). *N*-terminal sequencing of recombinantly expressed murine TMIE is also consistent with cleavage of a signal peptide (Supplementary Fig. 2), as are truncation experiments of zebrafish TMIE ^10^, supporting the hypothesis that in *C. elegans* the first ∼17 residues of TMIE functions as a signal peptide.

There is a striking intramembranous ‘cavity’ between TMIE and TMC-1 that is occupied by at least eight lipid molecules. Several lipids make hydrophobic contacts with nonpolar residues in TMIE and the putative pore-forming TMC-1 helices TM6 and TM8, bridging the two subunits. Consistent with the observed lipid density in the cryo-EM maps, molecular dynamics (MD) simulations independently identify multiple lipids in this cavity (Extended Data Fig. 9). Notably, C44 of TMIE on the cytosolic boundary of the transmembrane domain is palmitoylated, with the acyl chain extending along TMC-1 TM8 (Fig. 3d, e). The location of TMIE near the putative TMC-1 pore and its lipid interactions suggests roles for TMIE, and possibly lipids, in gating by sensing membrane tension. This idea is supported by recent studies in mouse cochlear hair cells, which demonstrated that TMIE binds to phospholipids and its association with lipids is important for TMC-1 MT ^11^.

### CALM-1 subunits cloak cytoplasmic surfaces of TMC-1

Calcium and integrin-binding protein 2 (CIB2) and its homolog, CIB3, modulate the activity of the MT complex and bind to the TMC-1 subunit ^12, 14^. In harmony with the role of CIB2/CIB3 in MT channel function, mutants of CIB2 are associated with non-syndromic hearing loss ^25, 36, 37^. Our MS results (Fig. 1d and Extended Data Fig. 2) demonstrate that CALM-1, the *C. elegans* ortholog of CIB2, co-purifies with TMC-1, consistent with CALM-1 residing within the TMC-1 complex. Inspection of the map of the TMC- 1 complex reveals density features for two CALM-1 subunits on the cytosolic faces of each TMC-1 protomer. By exploiting the crystal structure of CIB3 in complex with a TMC-1 peptide ^14^, we fit models of CALM-1 to their respective density features (Fig. 4a). Like other CIB proteins, CALM-1 has three EF-hand motifs, two of which are located proximal to the C-terminus and harbor clearly bound Ca^2+^ ions. Following superposition, the root-mean-square deviation (RMSD) between CALM-1 and CIB3 from the CIB3-TMC-1 peptide complex is 0.69 Å, and together with a substantial sequence similarity, underscore the conservation of sequence and structure between the worm and mouse proteins (Extended Data Fig. 8). Extensive interactions knit together CALM-1 and TMC-1, involving a buried surface area of ∼2,903 Å^2^, and suggesting that CALM-1 may bind to TMC-1 with high affinity (Fig. 4a). Three distinct regions of CALM-1 interact with cytosolic helical features of TMC-1, the first of which involves TMC-1 helices H1 to H3, oriented like ‘paddles’ nearly parallel to the membrane (Fig. 4b, c). Prominent interactions include side chains in the loop between H1 and H2, which form hydrophobic contacts with CALM-1, together with acidic residues on CALM-1 that create a negatively charged surface juxtaposed to a complementary positively charged surface on the H1-H3 paddle (Fig. 4c). The second binding interface is through a hydrophobic pocket of CALM-1, comprised of its EF-hand motifs, and the cytosolic H5-H6 helices of TMC-1 (Fig. 4d), reminiscent of the CIB3/TMC-1 peptide structure. Aliphatic and aromatic residues, including L308, F309 and Y314 of TMC-1 are docked into the conserved hydrophobic core in CALM-1, further stabilizing the complex by burial of substantial non-polar surface area. Lastly, amino acids D192, R195 and R200 at the C-terminus of CALM-1 interact with R780, D313 and E160 of TMC-1, respectively, forming conserved salt bridges through the buried short helix (191-197) of CALM-1 (Fig. 4e). This interface shows that CALM-1 directly engages with the transmembrane helices of TMC-1 via the loop between TM8 and TM9, and is thus positioned to modulate the ion channel function.

Multiple missense mutations of human CIB2 or TMC-1 are associated with non-syndromic hearing loss by either impeding the interaction between TMC-1 and CIB2 or by reducing the Ca^2+^ binding propensity of CIB2 ^14^. Several of these residues, E178D of human TMC-1 and E64D, F91S, Y115C, I123T, and R186W of CIB2, are structurally conserved in the *C. elegans* TMC-1 and CALM-1 complex (Extended Data Fig. 8). Our structure illuminates the proximity of the CALM-1 Ca^2+^ binding sites to the CALM-1 and TMC-1 interface, thus underscoring the roles of both Ca^2+^ and CALM-1 in sculpting the conformation of the TMC-1 and by extrapolation, providing a structural understanding of CIB2 in hair cell function.

MS analysis of the TMC-1 complex indicated the presence of a soluble protein known as ARRD-6. Upon classification of the single particle cryo-EM data, we noticed one, non 2-fold symmetric, 3D class defined by an elongated density feature protruding from the CALM-1 auxiliary subunit and we hypothesized that it could correspond to ARRD-6. Arrestins are composed of an N- and a C- domain, each comprised of β-sandwich motifs, which together give rise to a protein with an elongated, bean-like shape. We fit the predicted structure of ARRD-6 into the corresponding density feature and although the local resolution of the ARRD-6 region is lower than that of central region of the complex, the fit yielded overall correlation coefficients of 0.69 (mask) and 0.65 (volume) (Extended Data Fig. 5). Moreover, density features for the ARRD-6 β-sheets are clearly observed at the binding interface with CALM-1, as well as for the crossed elongated loops of the N- and C-domain at the central crest, further supporting the assignment of the density feature to ARRD-6. We observed a ‘C-edge loop’ structure, positioned at the distal edge of the β-strands in the C-domain, a feature which functions as a membrane anchor and is necessary for activation of arrestin (Fig. 4f, g)^38^. The C-edge loop of ARRD-6 includes W197 and multiple cysteine residues (Fig. 4g), the latter of which may be palmitoylated and thus poised for membrane anchoring ^39^. Additional contacts between CALM-1 and ARRD-6 involve a loop of CALM-1 (P51-K67) with the β-strands in the C-domain of ARRD-6 (Fig. 4h). At present, there is no known role for an arrestin in the function of the TMC channel of *C. elegans*, nor in the vertebrate TMC-1 complex. We speculate that ARRD-6 may play a regulatory role in TMC-1 channel function or it may be involved in endocytosis of the TMC-1 complex by recruitment of cytoskeleton proteins, akin to the role that α-arrestin plays in GPCR regulation ^40, 41^. At this juncture we do not know why we observed only a single ARRD-6 subunit bound to the complex, as there is sufficient space for two. Perhaps one subunit unbound from the complex or the second subunit is only partially occupied. Further experiments are required to address these questions.

### Mapping the putative ion channel pore

Single-channel currents measured from cochlear hair cells demonstrate that the mammalian MT complex is cation-selective with a high permeability for Ca^2+^ ^42^. TMC-1, or TMC-2, are likely the pore-forming subunits of the mammalian MT complex and cysteine mutagenesis experiments have pinpointed several pore-lining residues critical for TMC-1-mediated MT ^7, 8^. While *C. elegans* TMC-1 mediates mechanosensitivity in worm OLQ neurons and body wall muscles ^13^, its ion selectivity and permeation properties are not known, largely due to challenges associated with heterologous expression of the recombinant complex and with vanishingly small amounts of native material. Interestingly, *C. elegans* and murine TMC-1 also function as Na^+^ permeable leak channels that modulate the resting membrane potential via a depolarizing background leak conductance, suggesting that TMC-1 may serve multiple cellular roles ^23, 43^ and indicating that the channel pore is permeable to a greater diversity of ions than previously appreciated.

To gain insight into the nature and function of the *C. elegans* TMC-1 ion conduction pathway, we superimposed the TMC-1 subunit onto the structures of TMEM16A and OSCA1.2, revealing a similar architecture among the transmembrane domains (Extended Data Fig. 6). The TMEM16A and OSCA1.2 dimer assemblies harbor two pores, one within each subunit, that are defined by helices TM3-TM7.

Structural similarities between TMEM16a and OSCA1.2 suggest that TMC-1 may also have two pores and could conduct ions through a structurally analogous pathway composed of TM4-TM8 (Fig. 5a). The putative ion conduction pathway appears closed, with a narrow pore blocked by three constrictions (Fig. 5b, c). Polar and basic residues line the first constriction site near the extracellular pore entrance and nonpolar residues dominate the second constriction site, located ∼20 Å farther ‘down’ the conduction pathway, towards the cytoplasm. The remaining 40 Å of the conduction pathway is lined by mostly polar and charged residues (Fig. 5d). Seven basic residues line the pore, two of which (H404 and H731) partially define the third and narrowest constriction site. We visualized two spherical, non-protein densities near two acidic residues (D683 and D695) that may correspond to bound cations (Fig. 5b). At the present resolution, however, we cannot determine if these features are Ca^2+^ ions. Both Asp residues are conserved in human TMC-1, suggesting that they are important for ion coordination. While the ionizable residues lining the pore are predominately basic and thus not in keeping with a canonical Ca^2+^-permeable channel, which is typically dominated by acidic residues, the overall residue composition is similar to the mechanosensitive ion channel OSCA1.2 ^29^. OSCA1.2 displays stretch-activated non-selective cation currents with 17-21% Cl^-^ permeability ^44^, suggesting *C. elegans* TMC-1 may exhibit similar permeation properties.

To visualize the ion conduction pore of the vertebrate MT complex, we exploited the *C. elegans* structure and constructed a homology model of the human TMC-1 complex that includes TMC-1, CIB2 and TMIE (Supplementary Fig. 3). Upon inspection of the human structure, we found that the putative pore is lined by two basic residues and five acidic residues, in keeping with the channel being permeable to Ca^2+^. In addition, there are relatively more polar residues compared to the worm ortholog and the histidine residues that occlude the second constriction site in *C. elegans* TMC-1 are replaced by M418 and A579 in the human model. The vertebrate MT complex also endows hair cells with permeability to organic molecules, including the dye FM1-43 ^45, 46^. While our structure does not provide direct insight into the pathway of small molecule permeation, several hydrophobic crevices, including the lipid-lined space between TMC-1 and TMIE, provide possible routes for the transmembrane passage of small molecules such as FM1-43.

### ‘Expanded’ and ‘Contracted’ conformations

We discovered a second conformation of the TMC-1 complex via 3D classification, termed the ‘C’ conformation (Fig. 6a). The TMC-1 subunits in the ‘E’ and ‘C’ conformations have a similar structure and both have closed ion channels. In the ‘C’ conformation, however, the TM10 helix is bent ∼9° compared with that of ‘E’, and we can observe one more helical turn of TM10 in the ‘E’ conformation. Upon successively superimposing the TMC-1 subunits from the ‘C’ and ‘E’ complexes we observe that in the ‘E’ state, each half of the TMC-1 complex, composed of TMC-1/CALM-1/TMIE subunits, are rotated by ∼8° in comparison to the ‘C’ state, by way of an axis of rotation that is located near the TMC-1 H7-H8 helices and oriented approximately parallel to the membrane. The movement of each half of the complex, when viewed parallel to the membrane plane, thus resembles the motion of an accordion, with the cytoplasmic regions of the complex undergoing relatively larger conformational displacements in comparison to those on the extracellular side of the membrane. Indeed, in comparing the ‘C’ and ‘E’ states, the amphipathic TMC-1 H3 helices move farther apart by ∼11 Å, thus underscoring the magnitude of the conformational change. Together, these results illustrate the conformational plasticity of the TMC-1 complex and, reciprocally, the possibility that deformations of the membrane may induce conformational changes in the TMC-1 complex.

### Membrane embedding of the TMC-1 complex

To understand how the TMC-1 complex interacts with individual lipids as well as with the lipid bilayer, we performed all-atom (AA) and coarse-grained (CG) MD simulations on the complex embedded in a membrane composed of phospholipids and cholesterol (Extended Data Fig. 9a). The AA set included three independent simulation replicas yielding a collective sampling time of 3 μs, whereas the CG simulations were performed for 8 μs on a system including 4 TMC-1 complexes in a larger membrane patch resulting in a sampling time of 32 μs of lipid-protein interactions (Extended Data Fig. 9a). The equilibrated structure of the membrane around the TMC-1 complex indicates an unusually deep penetration and anchoring of the amphipathic, ‘paddle’ H3 helix into the cytosolic leaflet of the bilayer (Fig. 6c). In agreement with the cryo-EM density maps, the simulations show that phospholipids and cholesterol occupy the cavity between TMIE and TMC-1 and cholesterol is enriched in crevices near the 2-fold related, TM10 helices at the TMC-1 subunit interface, together supporting the importance of lipids in the structure and function of the complex (Extended Data Fig. 9b, c). The TMC-1 complex also distorts the membrane bilayer, promoting both thinning and thickening of the membrane, with especially prominent thinning of the cytoplasmic leaflet within the region of H3 helix insertion (Fig. 6d and Extended Data Fig. 9d). Strikingly, the effect of the TMC-1 complex on the membrane bilayer is long-range and propagates up to ∼50 Å from the protein (Extended Data Fig. 9d), thus suggesting an interplay between membrane structure and the function of the TMC-1 complex.

## Summary

The molecular structures of the TMC-1 complex reveal the identity, architecture, and membrane association of key subunits central to vertebrate and *C. elegans* mechanosensory transduction. The accordion-shaped, 2-fold symmetric complex harbors TMIE subunits poised like ‘handles’ perpendicular to the membrane, and amphipathic TMC-1 H3 helices inserted and parallel to the membrane plane, each providing possible mechanisms for direct or indirect transduction of force to ion channel gating, respectively. In vertebrates, protocadherin-15 transduces force to stereocilia tips, opening the MT channel. Prior studies suggest that protocadherin-15 forms a stable, dimeric complex with LHFPL5 yet also interacts with TMC-1 and TMIE subunits ^9, 15, 17, 47, 48^. How might protocadherin-15, either alone or in complex with LHFPL5, interact with the TMC-1 complex? One possibility is that the protocaderin-15 dimer is situated coincident with the 2-fold axis of the TMC-1 complex, with procadherin-15 TMs ‘surrounding’ the TMC- 1 TM10 helixes. This ‘closed’ symmetric dimeric complex would enable tension on protocadherin-15 to be directly transduced to the TMC-1 complex via the protocadherin-15 contacts with the TM10 helices. Alternatively, protocadherin-15 dimers could interact with TMIE helices, with one protocadherin subunit interacting with a single TMIE subunit, thus forming an ‘open’ complex in which the ‘unpaired’ protocadherin-15 subunit could interact with a TMIE subunit from another TMC-1 complex. This model not only provides a direct mechanism of force transduction from protocadherin-15 to TMIE and then to the TMC-1 ion channel pore, but it also provides a mechanism for the clustering of TMC-1 complexes ^49^. In additional to direct transduction of force, we also speculate that H3 of the TMC-1 subunit acts like a paddle in the membrane that will move ‘up’ or ‘down’ as the membrane thins or thickens, thus providing a mechanism for force coupling to the channel via the membrane. Further studies of open-channel conformations of the *C. elegans* TMC-1 complex, in addition to structures of the vertebrate MT complex, will be required to more fully elucidate the mechanisms of force transduction. Nevertheless, these TMC-1 complexes provide a framework for structure-based mechanisms of touch in *C. elegans* and of hearing and balance in vertebrates.

## Materials and Methods

### Transgenic worm design

The strain PHX2173 *tmc-1(syb2173)* was generated by SunyBiotech using CRISPR/Cas9 genome editing and is referred to as the *tmc-1::mVenus* line (Supplementary Fig. 1). The TMC-1-mVenus-3xFLAG sequence was inserted prior to the stop codon of the endogenous *tmc-1* gene (Wormbase: T13G4.3.1). The genotype was confirmed using PCR and primers ER02-seq-s (ATTAGATCCCGCAAGAGAAT) and ER02-seq-a (AAGGTGATATGAACGAACCG), which bind 452bp upstream and 408 bp downstream from the insertion site, respectively to amplify the region of interest. The PCR product was subsequently sequenced using primers ER02-mid-s (CATGAAGCAACACGACTTCT) and ER02-mid-a (TCTTCGATGTTGTGACGGAT), which bind within the TMC-1-mVenus-3xFLAG sequence. To enable elution of the engineered TMC-1 complex from affinity chromatography resin, a PreScission protease (3C) cleavage site was placed between the C-terminus of TMC-1 and the mVenus fluorophore.

### Spectral confocal imaging

Adult worms were immobilized in M9 buffer (22 mM KH2PO4, 42 mM Na2HPO4, 86 mM NaCl, and 1 mM MgCl2) containing 30 mM sodium azide and placed on slides that were prepared with ∼4 mm agar pads. Spectral images were acquired on a Zeiss 34-channel LSM 880 Fast Airyscan inverted microscope with a 40x 1.2 NA water-immersion objective lens. Linear unmixing was employed to distinguish between the mVenus signal and autofluorescence. The autofluorescence signal was subtracted from each image. The 3D z-stack information is presented in 2D after performing a maximum intensity projection.

### Large scale *C. elegans* culture

All *C. elegans* strains were maintained and grown according to Wormbook methods (http://www.wormbook.org). For large scale liquid culture, nematode growth medium (NGM) agar plates were prepared and spread with *E. coli* strain HB101, allowing the bacterial lawn to grow overnight at 37 °C. Worms were transferred to the NGM plates and grown for 3-4 days at 20 °C until HB101 cells were depleted. Worms on the plates were transferred to a liquid medium in 2L baffled flasks, supplemented with HB101 (∼15g per 500 mL medium) and streptomycin (50 μg/mL), and worms were grown at 20 °C with vigorous shaking (150 rpm) for 70-72 hours. To harvest worms, the liquid culture flasks were placed on ice for 1 hour to allow the worms to settle. The media was removed, and the worm slurry was collected in a tube, washed twice with 50 mL of ice cold M9 buffer by successive centrifugation (800 x g for 1 minute) and resuspension. Worms were ‘cleaned’ by sucrose density centrifugation at 1500 x g for 5 minutes after bringing the volume of worm slurry up to 25 mL with M9 buffer and adding 25 mL of ice cold 60% (w/v) sucrose. The worm layer on top was recovered and placed in a new tube and then washed twice with 50 mL of ice cold M9 buffer. The volume of the worm pellet was measured and the same volume of M9 buffer was added to the tube and worm balls were made by dripping the slurry into liquid nitrogen. The worm balls were stored at −80 °C until further use.

### Isolation of the native TMC-1 complex

Approximately 80 g of frozen worm balls were disrupted using a ball mill (MM400, Retch) where the grinding jar and ball were pre-cooled in liquid nitrogen. Disrupted worm powder was solubilized at 4 °C for 2 hours in a buffer containing 50 mM Tris-Cl (pH 9.3), 50 mM NaCl, 5 mM EDTA, 2% (w/v) glyco-diosgenin (GDN), and protease inhibitors (0.8 µM aprotinin, 2 µg/mL leupeptin and 2 µM pepstatin). After centrifugation at 40,000 rpm (186,000 x g) for 50 minutes, the supernatant was applied to anti-FLAG M2 affinity resin and incubated overnight on a rotator at 4°C. The resin was washed 5 times with a buffer containing 20 mM Tris-Cl (pH 8.5), 150 mM NaCl and 0.02% (w/v) GDN, using a volume of buffer that was 200-fold the volume of the resin. The TMC-1 complex was eluted by incubating with 40 μg of 3C protease at 4 °C for 4 hours on the rotator. Subsequently, the solution was supplemented with 3 mM CaCl2, final concentration, and the eluate was filtered with a 0.22 µm centrifuge tube filter. The concentrate was loaded onto a size-exclusion chromatography (SEC) column (Superose 6 Increase 10/30 GL, GE Healthcare), equilibrated in a buffer composed of 20 mM Tris-Cl (pH 8.5), 150 mM NaCl, 0.02% (w/v) GDN and 3 mM CaCl2. The peak fractions from the putative dimeric TMC-1 complex were pooled and concentrated for cryo-EM grid preparation. Approximately 50 ng of TMC-1 was isolated from 80 g of worm balls, which translates to approximately 6 × 10^7^ worms. The amount of protein was determined via mVenus fluorescence based on a standard plot. The estimated total amount of the TMC-1 complex including TMC-1, CALM-1 and TMIE is 60 ng. The isolated native TMC-1 sample was analyzed by SDS-PAGE (sodium dodecyl sulfate–polyacrylamide gel electrophoresis) and the protein bands were visualized by silver staining. For mass-spectrometry analysis, the putative dimeric TMC-1 complex peak was pooled and concentrated to a volume of 50 µL for further use. The same isolation method was utilized to make the wild-type worm sample from the *C. elegans* N2 strain for use as a control in the mass spectrometry experiments in order to evaluate non-specific binding of *C. elegans* proteins to anti-FLAG M2 affinity resin.

### Isolation of the native TMC-1 complex for SiMPull

The native TMC-1 complex, bound to anti-FLAG M2 affinity resin, was eluted via a buffer composed of 20 mM Tris-Cl (pH 8.5), 150 mM NaCl and 0.02% (w/v) GDN, supplemented with 1 mg/mL 2X FLAG peptide at 4 °C for 40 minutes on a rotator. The eluate was concentrated and subjected to further purification on a SEC column. The putative dimeric TMC- 1 complex peak was pooled and used for SiMPull.

### SiMPull

Coverslips and glass slides were cleaned, passivated and coated with a solution consisting of 50 mM methoxy polyethylene glycol (mPEG) and 1.25 mM biotinylated PEG in water. A flow chamber was created by drilling 0.75 mm holes in a quartz slide and by placing double-sided tape between the holes. A coverslip was placed on top of the slide and the edges were sealed with epoxy, creating small flow chambers. A solution of phosphate buffered saline (PBS) that included 0.25 mg/mL streptavidin was then applied to the slide, allowed to incubate for 5 minutes, and washed off with a buffer consisting of 50 mM Tris, 50 mM NaCl and 0.25 mg/mL bovine serum albumin (BSA), pH 8.0 (T50 BSA buffer). Biotinylated anti-GFP nanobody in T50 BSA at 10 µg/mL was applied to the slide, allowed to incubate for 10 minutes, and washed off with 30 µL buffer A (20 mM Tris, pH 8.0, 150 mM NaCl, 0.02% (w/v) GDN, 3 mM CaCl2).

The TMC-1 complex was isolated as previously described under ‘isolation of the native TMC-1 complex for SiMPull’. The complex was purified by SEC, diluted 1:200, and immediately applied to the chamber. After a 5-minute incubation, the slide was washed with 30 μL buffer A and the chamber was imaged using a Leica DMi8 TIRF microscope with an oil-immersion 100x objective. Images were captured using a back-illuminated EMCCD camera (Andor iXon Ultra 888) with a 133 × 133 µm imaging area and a 13 µm pixel size. This 13 µm pixel size corresponds to 130 nm on the sample due to the 100x objective. To estimate non-specific binding to the glass slide, the purified TMC-1 complex was applied to a separate chamber wherein the anti-GFP nanobody was not included and the other steps remained identical. The observed spot count from this chamber was used to estimate the number of background fluorescence spots.

Photobleaching movies were acquired by exposing the imaging area for 60 seconds. To count the number of TMC-1 subunits, single-molecule fluorescence time traces of the mVenus-tagged TMC-1 complex were generated using a custom python script. Each trace was manually scored as having one to three bleaching steps or was discarded if no clean bleaching steps could be identified. The resulting distribution of bleaching steps closely matches a binomial distribution for a dimeric protein based on an estimated GFP maturation of 80%. A total of 600 molecules were evaluated from three separate movies. Scoring was verified by assessing the intensity of the spot; on average, the molecules that bleach in 2 steps were twice as bright as those that bleach in 1 step.

### Mass spectrometry

The purified TMC-1 complex sample was dried, dissolved in 5% sodium dodecyl sulfate, 8 M urea, 100 mM glycine (pH 7.55), reduced with (tris(2-carboxyethyl)phosphine (TCEP) at 37 °C for 15 min, alkylated with methyl methanethiosulfonate for 15 min at room temperature followed by addition of acidified 90% methanol and 100 mM triethylammonium bicarbonate buffer (TEAB; pH 7.55). The sample was then digested in an S-trap micro column briefly with 2 µg of a Tryp/LysC protease mixture, followed by a wash and 2 hr digestion at 47 °C with trypsin. The peptides were eluted with 50 mM TEAB and 50% acetonitrile, 0.2% formic acid, pooled and dried. Each sample was dissolved in 20 µL of 5% formic acid and injected into Thermo Fisher QExactive HF mass spectrometer. Protein digests were separated using liquid chromatography with a Dionex RSLC UHPLC system, then delivered to a QExactive HF (Thermo Fisher) using electrospray ionization with a Nano Flex Ion Spray Source (Thermo Fisher) fitted with a 20um stainless steel nano-bore emitter spray tip and 1.0 kV source voltage. Xcalibur version 4.0 was used to control the system. Samples were applied at 10 µL/min to a Symmetry C18 trap cartridge (Waters) for 10 min, then switched onto a 75 µm x 250 mm NanoAcquity BEH 130 C18 column with 1.7 µm particles (Waters) using mobile phases water (A) and acetonitrile (B) containing 0.1% formic acid, 7.5- 30% acetonitrile gradient over 60 min and 300 nL/min flow rate. Survey mass spectra were acquired over m/z 375-1400 at 120,000 resolution (m/z 200) and data-dependent acquisition selected the top 10 most abundant precursor ions for tandem mass spectrometry by higher energy collisional dissociation using an isolation width of 1.2 m/z, normalized collision energy of 30 and a resolution of 30,000. Dynamic exclusion was set to auto, charge state for MS/MS +2 to +7, maximum ion time 100 ms, minimum AGC target of 3 × 10^6^ in MS1 mode and 5 × 10^3^ in MS2 mode. Data analysis was performed using Comet (v. 2016.01, rev.^50^ against a January 2022 version of canonical FASTA protein database containing *C. elegans* uniprot sequences and concatenated sequence-reversed entries to estimate error thresholds and 179 common contaminant sequences and their reversed forms. Comet searches for all samples performed with trypsin enzyme specificity with monoisotopic parent ion mass tolerance set to 1.25 Da and monoisotopic fragment ion mass tolerance set at 1.0005 Da. A static modification of +45.9877 Da was added to all cysteine residues and a variable modification of +15.9949 Da on methionine residues. A linear discriminant transformation was used to improve the identification sensitivity from the Comet analysis ^51, 52^. Separate histograms were created for matches to forward sequences and for matches to reversed sequences for all peptides of seven amino acids or longer. The score histograms of reversed matches were used to estimate peptide false discovery rates (FDR) and set score thresholds for each peptide class. The overall protein FDR was 1.2%.

### Cryo-EM sample preparation

A volume of 3.5 μL of the concentrated TMC-1 complex was applied to a Quantifoil grid (R2/1 300 gold mesh, covered by 2 nm continuous carbon film), which was glow-discharged at 15 mA for 30 seconds in the presence of amylamine. The grids were blotted and flash frozen using a Vitrobot mark IV for 2.5 seconds with 0 blot force after 30 seconds wait time under 100% humidity at 15 °C. The grids were plunge-frozen into liquid ethane, cooled by liquid nitrogen.

### Data acquisition

The native TMC-1 complex dataset was collected on a 300 keV FEI Titan Krios microscope equipped with a K3 detector. The micrographs were acquired in super-resolution mode (0.4195 Å/pixel) with a magnification of 105kx corresponding to a physical pixel size of 0.839 Å/pixel. Images were collected by a 3×3 multi-hole per stage shift and a 6 multi-shot per hole method using Serial EM, with a defocus range of −1.0 to −2.4 µm. Each movie stack was exposed for 3.3 seconds and consisted of 50 frames per movie, with a total dose of 50 e^−^/Å^2^. A total of 26,055 movies were collected.

### Image processing

Beam-induced motion was corrected by patch motion correction with an output Fourier cropping factor of 1/2 (0.839 Å/pixel). Contrast transfer function (CTF) parameters were estimated by patch CTF estimation in CryoSparc v3.3.1 ^53^. A total of 25,852 movies were selected by manual curation and the particles were picked by using blob-picker with minimum and maximum particle diameters of 140 Å and 200 Å, respectively. Initially, 7.9 million particles were picked and extracted with a box size of 400 pixels and binned 4x (3.356 Å/pixel). After one round of 2D classification, ‘junk’ particles were removed, resulting in 3.2 million particles in total. The particles with the highest resolution features, approximately 1.5 million, were used for *ab initio* reconstruction. The full particle stack consisting of 3.2 million particles from 2D classification were then subjected to heterogeneous refinement using the reconstructed models from the *ab initio* reconstruction. Probable monomeric TMC-1 complexes, detergent micelles, and additional junk particles were removed in this step, yielding 1.65 million particles. Particles were then re-extracted from unbinned images. Subsequently, heterogeneous refinement using C1 symmetry was performed with the re-extracted 1.65 million particles, yielding 8 classes. Among them, three good classes composed of 667k particles were selected and used for further analysis. After one round of heterogeneous refinement with 4 classes in C2 symmetry, two classes containing 208k and 199k particles were discerned, each with distinct features and that we describe as the ‘contracted’ and ‘expanded’ forms, respectively. One more round of heterogeneous refinement was performed for both particle stacks to sort out groups of homogeneous particles from each class. To attain higher resolution and improved map quality, non-uniform refinement including defocus and global CTF refinement was performed in Cryosparc v3.3.1 of each individual class, with particle stack sizes of 141k (contracted) and 142k (expanded), resulting in resolutions at 3.09 Å and 3.10 Å, respectively.

Among the initial 8 classes from the heterogeneous refinement of 1.65 million particles, one of the classes, which contained 272k particles, had an additional density feature, proximal to CALM-1. Further heterogeneous refinement and 3D classification without alignment was carried out with this class to sort out heterogeneous particles. One more round of heterogeneous refinement in Cryosparc resulted in one promising particle class, containing 99k particles, out of four total classes. Non uniform refinement, including defocus and global CTF refinement, was performed with the selected class, resulting in a map at 3.54 Å resolution. To improve the density of unknown protein bound to CALM-1, local refinement in Cryosparc was performed using a mask, covering the ‘extra density’ and CALM-1.

### Structure determination and model building

The initial EM density map was sharpened with Phenix AutoSharpen ^54^, and both sharpened and unsharpened maps were used for structure determination. Various strategies including *de novo* building, structure prediction, docking and homologous modeling were used for model building. The transmembrane helices of TMC-1 (TM1–TM9, excluding TM10), predicted by Alphafold2 ^55^ as a template, were fit into the map with rigid body fitting in UCSF Chimera ^56^ and *de novo* model building using Coot ^57^. The possible ion permeation pore of the channel was determined by MOLE 2.0 ^58^. Carbohydrate groups were modeled to protruding densities of N209 on TMC-1, at a predicted *N*-linked glycosylation site.

To build the structure of CALM-1 into the ‘expanded’ conformation density map of the TMC-1 complex, we exploited the previously determined structure of CIB3 in complex with a TMC-1 peptide (PDB 6WUD). We docked CIB3 into the density map using rigid body fitting in UCSF Chimera, using the highly conserved H5-H6 helices of TMC-1 as a guidepost, and proceeded by introducing the sequence of CALM-1 into the model, followed by manual adjustment of the model using Coot. Conserved bulky side chains, including F84, Y129, and F197, that protrude into hydrophobic cavities and are facing the helices of TMC-1, facilitated the definition of the correct register of the CALM-1 sequence.

The auxiliary subunit, TMIE, was built manually into the density map of the ‘expanded’ conformation using Coot. The bulky side chain density of tryptophan (W25) and lipid modification on cysteine residue (C44) helped to assign the sequence register in the context of the density map. The model was refined against the sharpened map by real-space refinement in Phenix.

The following regions of TMC-1 were not modeled into the map because of weak or absent densities: The N-terminal region of TMC-1 (M1 to P73), the predicted loop region between TM5 and TM6 (S460 to N663) and the C-terminal region (L886 to D1285). The side chains with weak density on H1 (75-87) and TM10 (870-885) helices were modeled as alanine residues. The N-terminal region of CALM-1, from residues 1 to 17, and the amino acids of TMIE, including 1-17 and 64-117, were not modeled due to a lack of density. As discussed in the main text, we suggest that residues 1-17 of TMIE comprise a signal peptide.

For the modeling of the unknown density on CALM-1 we speculated that ARRD-6 was a possible candidate auxiliary protein based on the mass spectrometry results. Although the overall map quality of the putative ARRD-6 region was not sufficient for *de novo* model building, we could find several β-sheets with side chain densities on the map. Using the predicted structure of ARRD-6 and the crossed-protrusion of two loops of the N- and C- domains of arrestin (82-85 of the N-, and 249-256 of the C- domain), we could align the predicted ARRD-6 model into the unknown density, thus providing further evidence that the unknown density is ARRD-6. The estimated local resolution of ARRD-6 density ranges between 4-7 Å and the calculated Q-score of ARRD-6 model-to-map from MapQ ^59^ plugin in Chimera is 0.25, which corresponds to the estimated resolution of 4.91 Å, suggesting that the model is reasonably placed in the map. The final CC of the ARRD-6 and overall model are 0.42 and 0.69, respectively.

### Molecular Dynamics Simulations

The molecular dynamics (MD) simulations were performed on the ‘E’ conformation and at two different resolutions, coarse-grained (CG) and all-atom (AA). Starting from the cryo-EM modeled structure, a C-terminal carboxylic cap group, an N-terminal ammonium capping group, missing side chains and all the hydrogen atoms were modeled using the PSFGEN plugin of VMD ^60^. PROPKA was employed to estimate the pKa of titratable residues ^61, 62^. The modeled structure was then used for setting up the CG and AA simulations.

### Coarse-grained simulation setup

The Martini-based CG model ^63–65^ of the ‘E’ conformation was generated, employing the Martinize protocol as described in the Martini website (http://www.cgmartini.nl/), followed by applying an elastic network on atom pairs within a 10 Å cut-off. The CG parameters for the palmitoylated Cys in TMIE was obtained from a previous work ^66^. The initial orientation of the protein in the membrane was adopted from the Orientations of Proteins in Membranes (OPM) database. The protein complex was then inserted in a lipid bilayer composed of palmitoyl-oleoyl-phosphatidyl-ethanolamine (PE), palmitoyl-oleoyl-phosphatidyl-choline (PC), sphingomyelin (SM), and cholesterol with a molar ratio of 54:32:8:6. The secondary structure of the protein was derived from the AA model and maintained throughout the CG simulations. To enhance the sampling and improve statistics, four copies of the protein were embedded in a large patch (400 × 400 Å^2^) of lipid bilayer at an inter-protein distance of 200 Å, using the computer program ‘insane’ ^67^. The system was then solvated and ionized with 150 mM NaCl concentration employing insane (system size: 330k CG beads).

### All-atom simulation setup

The CG equilibrated protein-membrane complex at the end of the 8 μs CG simulation was back-mapped to a CHARMM-based AA model employing CHARMM-GUI ^68, 69^. Thus, one of the four replicas (a protein copy with membrane padding of approximately 40 Å) was isolated from the large membrane patch. DOWSER was used to internally hydrate the protein ^70, 71^. The protein-membrane system was then solvated with water including 150 mM NaCl in VMD (system size: 340k atoms). To improve the statics and further reduce any bias from the initial lipid placement, three independent membrane systems, with independently placed initial lipids, were generated using the Membrane Mixer Plugin (MMP) ^72^.

### Coarse-grained simulation protocol

CG systems were simulated using GROMACS ^73^, with the standard Martini v2.2 simulation parameters ^65^. The simulation was conducted with a 20 fs timestep. The temperature was fixed at 310 K using velocity-rescaling thermostat ^74^ with a time constant for coupling of 1 ps. A semi-isotropic 1-bar pressure was maintained employing the Berendsen barostat ^75^ with a compressibility of 3×10^−4^ bar and a relaxation time constant of 5 ps. The system was initially energy minimized for 1000 steps, followed by relaxation runs of 18 ns, while the lipid bilayer headgroups and protein backbones were restrained harmonically. During the initial 18 ns, the restraints applied on bilayer headgroups were removed stepwise (from *k* = 200 kJ.mol^-1^.nm^-2^ to zero), while the restraints on protein backbone (*k* = 1000 kJ.mol^-1^.nm^-2^) were unchanged. The four-protein system was then simulated for 8 μs, with restraints only applied to the protein backbones, resulting in a cumulative sampling of 32 μs (4 copies × 8 μs).

### All-atom simulation protocol

The AA converted system was simulated using the following protocol: (1) 5,000 steps of minimization, followed by 5 ns of relaxation, during which the proteins’ heavy atoms as well as the bound Ca^2+^ ions were harmonically restrained (*k* = 10 kcal.mol^-1^.Å^-2^) to their position in the cryo-EM model; (2) 1 ns of equilibration with harmonic restraints only on the protein backbone heavy atoms (*k* = 10 kcal.mol^−1^.Å^−2^). The coordination of Ca^2+^ ions in this step was maintained by the application of the Extra Bonds algorithm in NAMD ^76, 77^. (3) 200 ps of equilibration during which the restraints on the backbone were maintained whereas the Extra Bonds on the Ca^2+^ ions were removed. (4) Two additional replicas were generated employing the MMP plugin and 1 μs of production run was performed on each of the three replicas while only the protein backbone heavy atoms were restrained. Steps 1-3 were performed using NAMD2 ^76, 77^. The 1-μs production runs for all three replicas were conducted on Anton2 ^78^.

All AA simulations were performed using the fully atomistic CHARMM36m ^79^ and CHARMM36 ^80^ force fields for the protein and lipids, respectively. Water molecules were modeled with TIP3P ^81^. In NAMD simulations, a 12 Å cutoff was used for short-range, non-bonded interactions, with switching distance starting at 10 Å. Particle mesh Ewald (PME) was used to calculate long-range electrostatic interactions ^82^ with a grid density of 1 Å^−1^, and a PME interpolation order of 6. The SHAKE algorithm was used to constrain bonds involving hydrogen atoms ^83^. Temperature was kept constant at 310 K using Langevin thermostat with a damping coefficient of 1.0 ps^−1^. Pressure was maintained at 1 atm employing the Nosé-Hoover Langevin piston barostat with period and decay of 100 and 50 fs, respectively ^84, 85^. All systems were simulated in a flexible cell allowing the dimensions of the periodic cell to change independently while keeping the aspect ratio in the *xy* plane (membrane plane) constant. The timestep was set to 2 fs, and the PME and Lennard-Jones forces were updated at every other and each timesteps, respectively.

For Anton2 simulations, 310 K temperature and 1 bar pressure were kept by the Nosé–Hoover chain coupling and Martyna–Tuckerman–Klein schemes ^84^, as implemented using a multigrator scheme ^86^. M-SHAKE was used to constrain all the bonds to hydrogen atoms ^87^, and a 2.5 fs timestep was used in all the simulations. The long-range electrostatic interactions were calculated by employing the Fast Fourier Transform (FFT) method on Anton2 ^78^.

### Membrane thickness and lipid distribution analysis

The change in the thickness of each membrane leaflet in response to the protein was quantified in both CG and AA simulations by monitoring the *z* (membrane normal) distance of the phosphate groups of phospholipids with respect to the bilayer midplane, over the second half of each trajectory (last 4 μs of the CG simulations and the last 500 ns of the AA simulations). The thickness values were plotted using a histogram with 2 × 2 Å^2^ bins in the *xy* plane (membrane plane), for each leaflet individually. Cholesterol and phospholipid distributions were similarly calculated by histogramming the positions of the hydroxy (for cholesterol) and phosphate (for phospholipids) beads over the last 4 μs of the trajectory.

### Lipid depletion/enrichment analysis

First, individual lipid counts for all lipid species within 7 Å (using cholesterol or phospholipid phosphate beads) of the 4 protein copies over the 8 μs of the CG simulation were determined. A depletion/enrichment index for lipid type L was then defined using the following equation ^88^:

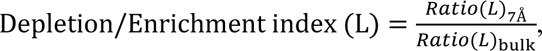

Where:

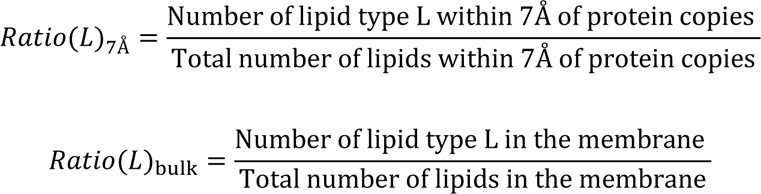

### Homology modeling of human TMC-1 complex

The cryo-EM structure of the ‘E’ conformation of *C. elegans* TMC-1 complex (containing 6 different chains: 2 TMC-1, 2 CALM-1 and 2 TMIE) was used as a template to build a homology model of human TMC-1 complex. Each chain in the template structure was isolated and its sequence was aligned to the corresponding human sequence with AlignMe ^89^. The aligned sequences were then used in the multi-chain capability of MODELLER ^90^ to generate a human TMC-1 complex. The discrete optimized protein energy (DOPE) ^91^ and GA341 ^92, 93^ methods were used to assess the quality of the generated model. The optimization was performed with a maximum iteration of 300 and the model with the best molecular probability density function (molpdf) was selected (Extended Data Fig. 9). The entire optimization cycle was repeated twice to obtain a better structure.

### Data Availability

The coordinates and volumes for the cryo-EM data have been deposited in the Electron Microscopy Data Bank under accession codes EMD-26741 (Expanded), EMD-26742 (Contracted), and EMD-26743 (with ARRD-6). The coordinates have been deposited in the Protein Data Bank under accession codes 7USW (Expanded), 7USX (Contracted), and 7USY (with ARRD-6).

## Acknowledgements

We thank T. Nicolson, P. Barr-Gillespie, D. Farrens, M. Mayer, A. Aballay, J. Ge, J. Elferich, and Gouaux and Baconguis lab members for helpful discussions, T. Provitola for assistance with figures, A. Reddy for mass spectrometric analysis, J. Meyers and S. Yang for help with cryo-EM screening and data collection, A. Chinn for help with worm growth, and R. Hallford for proof reading. Initial cryo-EM grids were screened at the Pacific Northwest Cryo-EM Center (PNCC), which is supported by NIH grant U24GM129547 and performed at the PNCC at OHSU, accessed through EMSL (grid.436923.9), a DOE Office of Science User Facility sponsored by the Office of Biological and Environmental Research. The large single particle cryo-EM data set was collected at the Janelia Research Campus of the Howard Hughes Medical Institute (HHMI). The OHSU Proteomics Shared Resource is partially supported by NIH core grants P30EY010572 and P30CA069533. This work was supported by NIH grant 1F32DC017894 to S.C. E.G. is an investigator of the HHMI. The simulations were supported by the NIH grants, P41-GM104601 and R01-GM123455 to E.T. Simulations were performed using allocations on Anton at Pittsburgh Supercomputing Center (award MCB100017P to E.T.), and XSEDE resources provided by the National Science Foundation Supercomputing Centers (XSEDE grant number MCA06N060 to E.T.).

## Author Contributions

H.J., S.C., and A.G. performed the experiments. H.J., S.C., and A.G., together with E.G., designed the project and wrote the manuscript. S.D.-G., A.R., and E.T. performed and analyzed MD simulations. All authors contributed to manuscript preparation.

## Additional Information

Supplementary Information is available for this paper.

## Materials and Correspondence

Correspondence and requests for materials should be addressed to E.G. Reprints and permissions information is available at www.nature.com/reprints.

---

**Extended Data Fig. 1.**
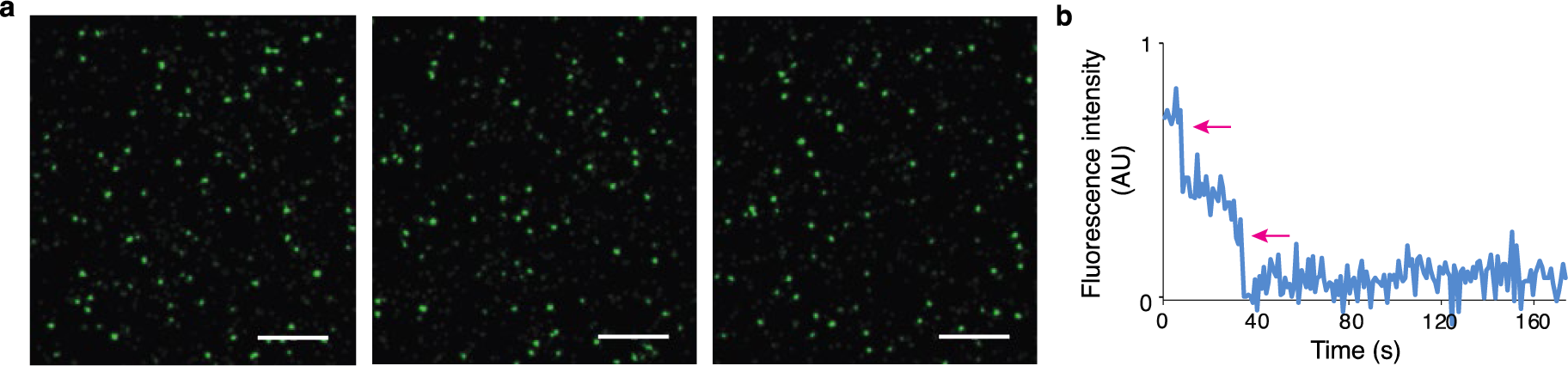
Representative TIRF images and photobleaching traces for the native, mVenus-tagged TMC-1 complex. **a,** Images are shown for the SEC-purified mVenus-tagged TMC-1 complex captured with biotinylated anti- GFP nanobody. Scale bar = 5 µm. **b,** Representative trace showing the two-step photobleaching (red arrows) of the mVenus-tagged TMC-1 complex.

**Extended Data Fig. 2.**
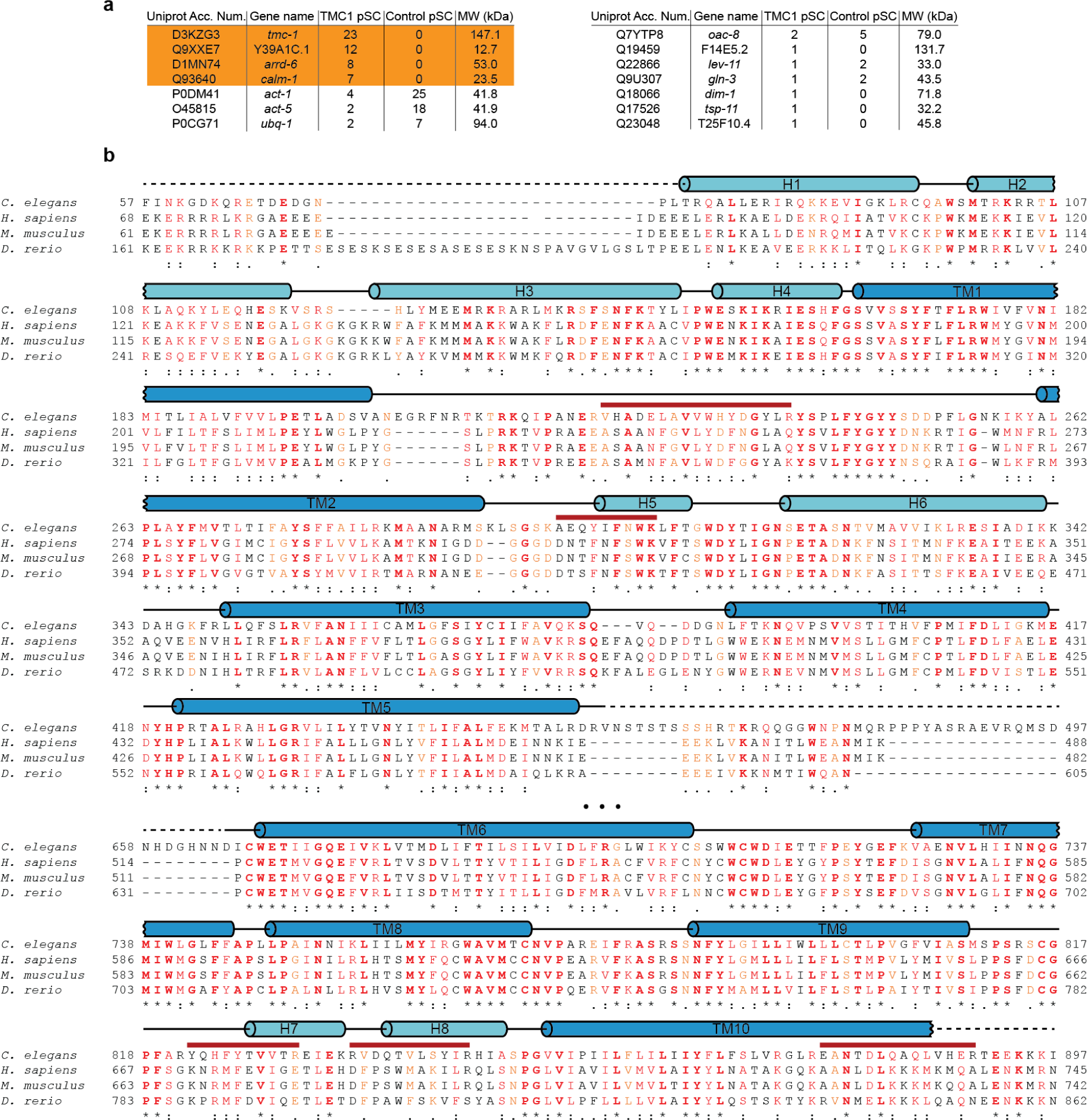
MS analysis of the TMC-1 complex. **a**, Proteins detected by MS, via their associated peptide fragments, are listed with their gene name and molecular mass. The number of identified unique peptides from both the native TMC-1 complex and from wild-type worms (*C. elegans* N2), used as a control, are also indicated. **b**, Amino acid sequence and secondary structure of *C. elegans* TMC-1 are shown. The secondary structure based on the cryo-EM structure is indicated above the sequences as boxes (α-helices), black lines (loop regions), or dashed lines (disordered residues). Peptides found by MS are indicated below the sequence (red lines). Note that the TMC-1 segments, corresponding to the sequence of 13-33, 557-566, 567-587, 877-890, 897-904, 917-927, 972-996, 1041-1052, 1177-1190, 1192-1216, and 1261-1269 are also found by MS, but not indicated in **b**.

**Extended Data Fig. 3.**
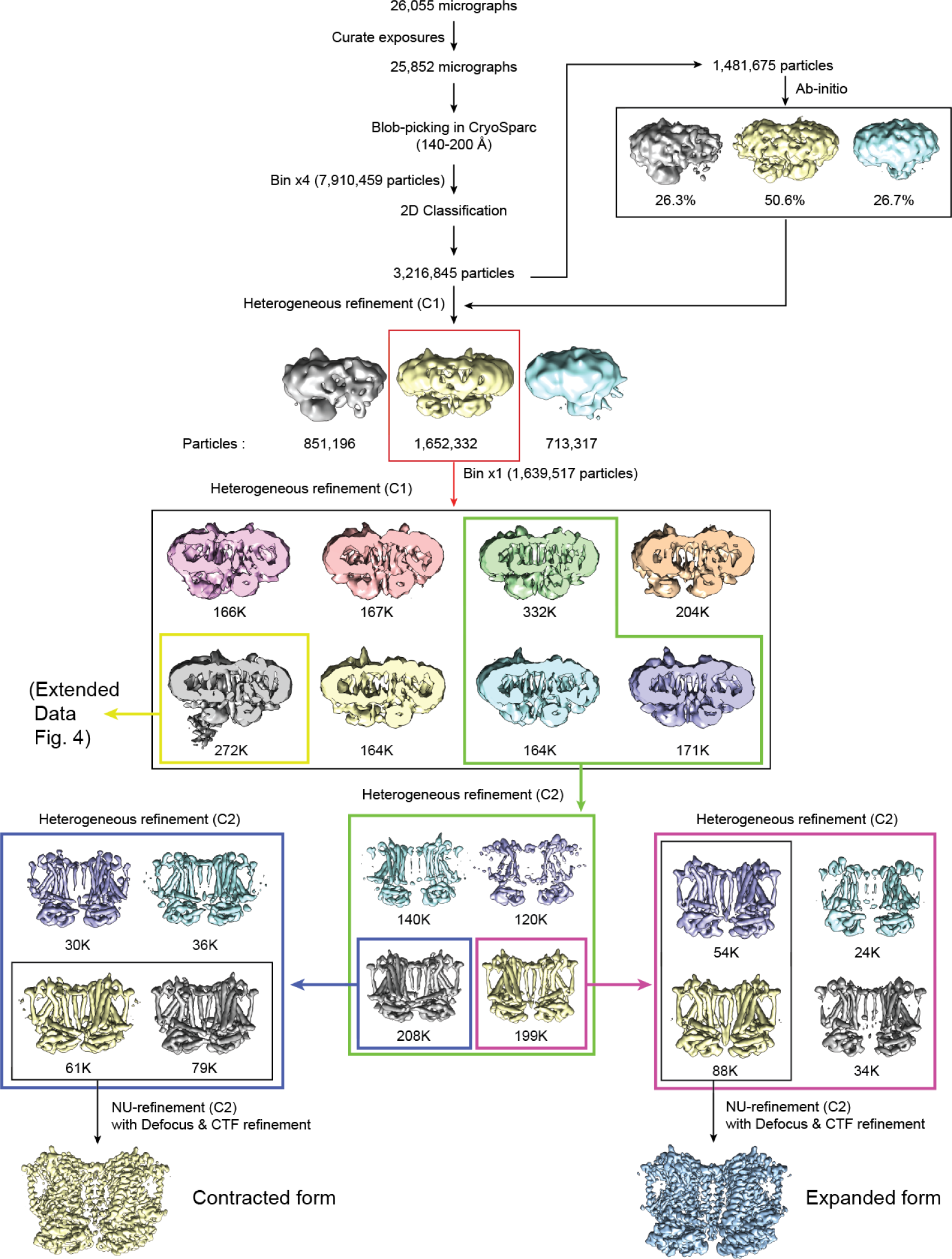
Cryo-EM processing workflow of ‘E’ and ‘C’ conformations. Flow chart for cryo-EM data analysis of ‘E’ and ‘C’ conformation of the TMC-1 complex.

**Extended Data Fig. 4.**
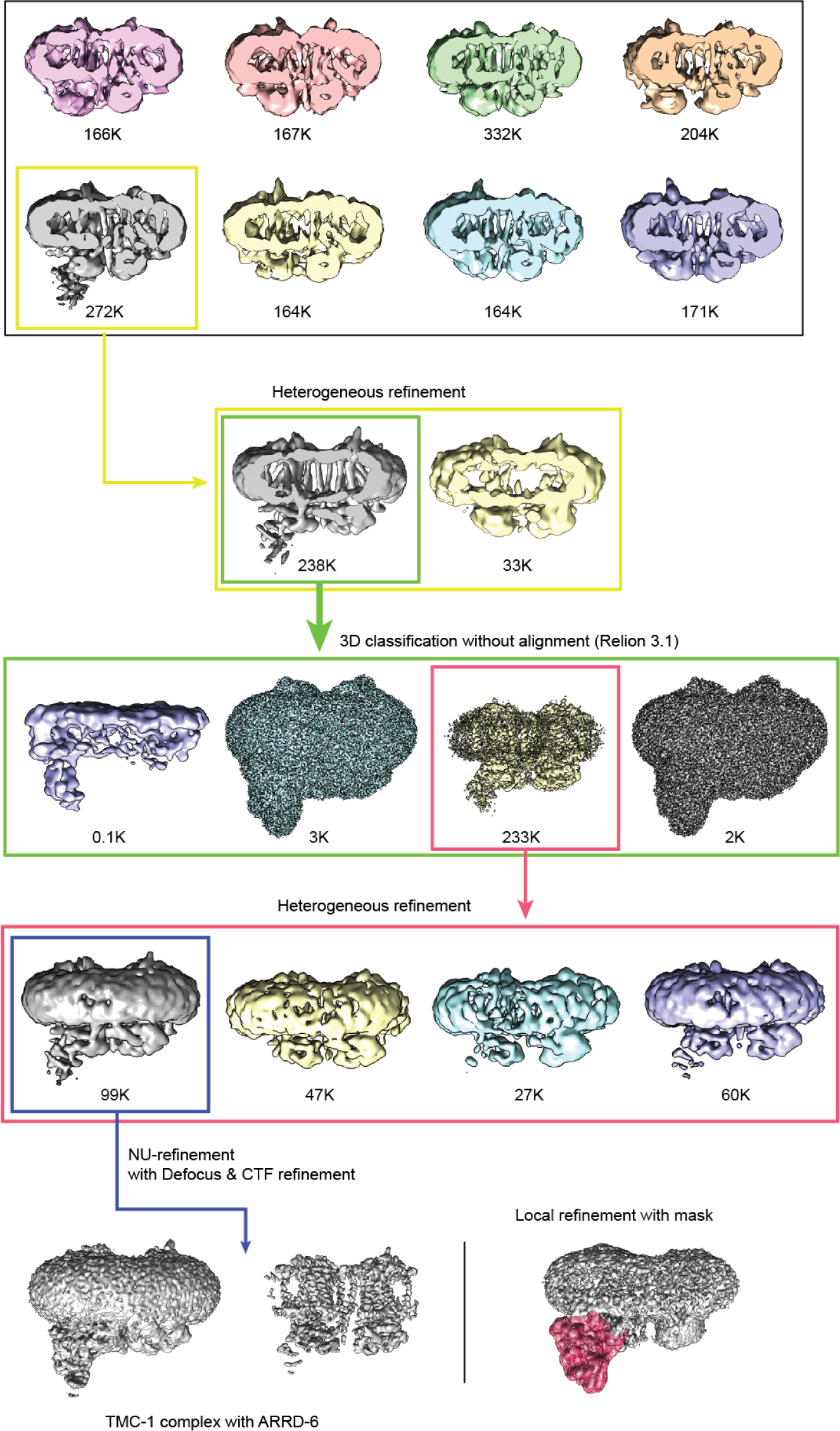
Cryo-EM processing workflow of TMC-1 complex with ARRD-6. Flow chart for cryo-EM data analysis of the TMC-1 complex with ARRD-6.

**Extended Data Fig. 5.**
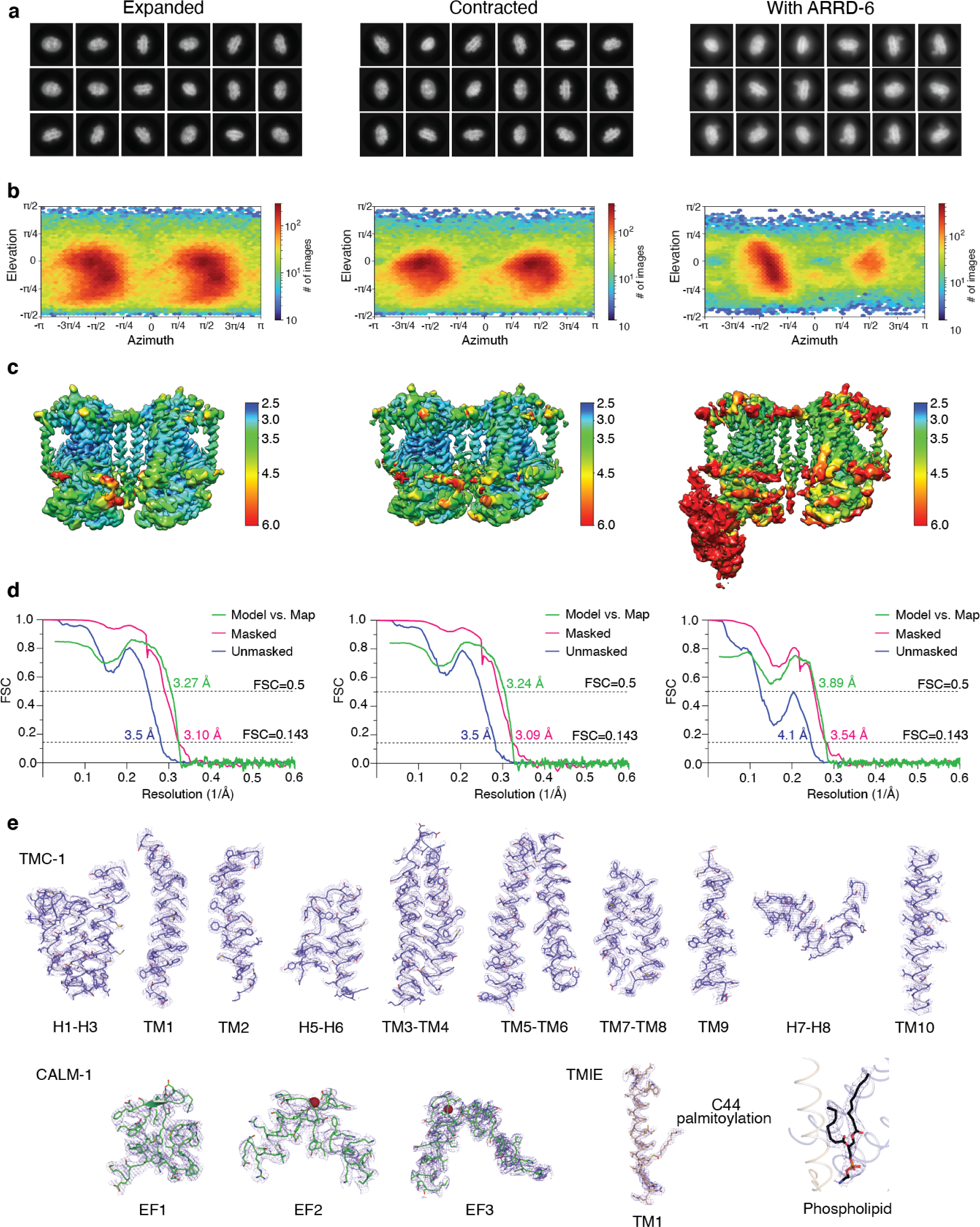
Cryo-EM classes, statistics, angular distributions and selected sections of density maps. **a**, Selected 2D class averages of ‘E’ and ‘C’, and with ARRD-6, respectively. **b**, Angular distributions of final reconstructions. **c**, Electron density map of each model colored by local resolution values. **d**, Fourier Shell Correlations (FSC) curve for each model. **e**, Fragments of cryo-EM density map and atomic model of each auxiliary subunit. The cryo-EM maps are shown as purple mesh.

**Extended Data Fig. 6.**
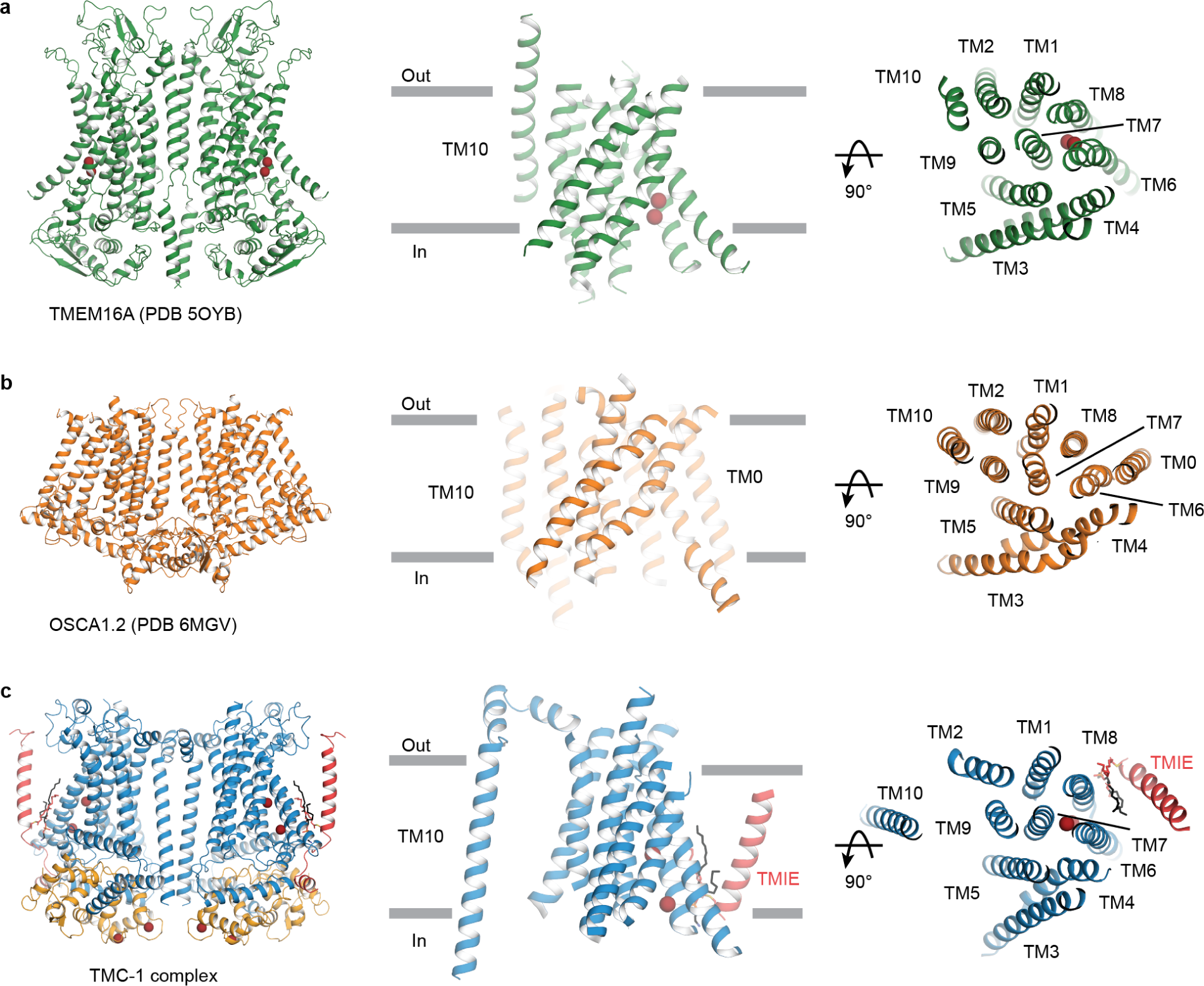
Structural comparison of TMEM16, OSCA1.2, and the TMC-1 complex. **a, b, c.** Structures of TMEM16A (5OYB), OSCA1.2 (6MGV), and the TMC-1 complex viewed from the same relative perspective. The side view of the transmembrane regions and the top-down views are shown in the cartoon model. Putative ions are shown as red spheres.

**Extended Data Fig. 7.**
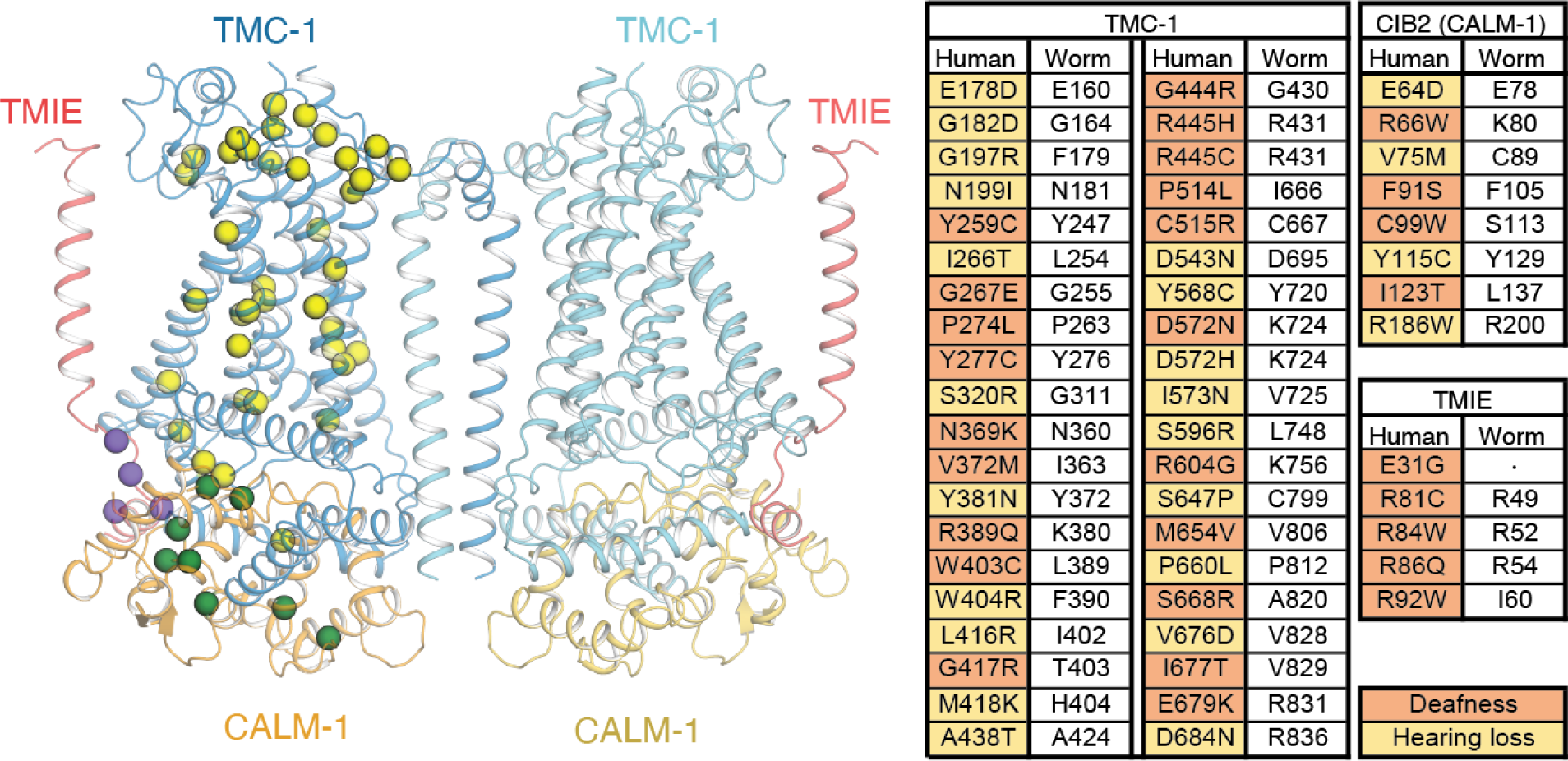
The locations of key mutations mapped to one protomer of the TMC-1 complex. The structure of the TMC-1 complex and the locations of mutations causing either hearing loss or deafness. Cα positions of the residues in question are shown as yellow (TMC-1), green (CALM-1), or purple (TMIE) spheres.

**Extended Data Fig. 8.**
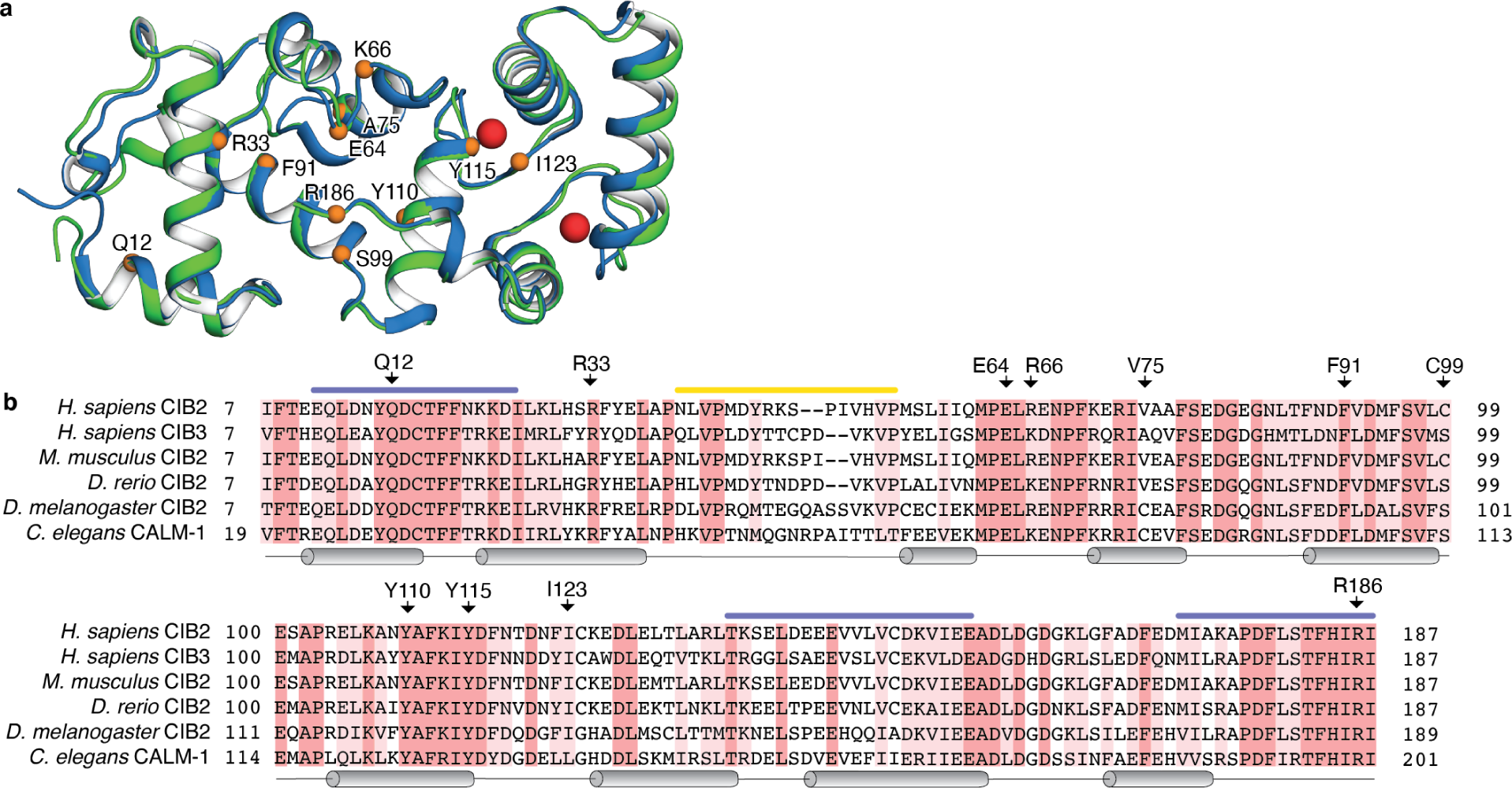
Structural and sequence alignment of CALM-1, CIB2, and CIB3. **a.** Superposition of *C. elegans* CALM-1 (green) and human CIB3 (blue, PDB 6WUD) using backbone α - carbon atoms highlights structural conservation (RMSD = 0.7 Å). Calcium (CALM-1) and magnesium (CIB3) ions are shown as red spheres. CIB2 residues implicated in deafness or hearing loss mutations are shown as orange spheres. **b.** Sequence alignment of human CIB2, human CIB3, and CIB2 orthologs from mouse, zebrafish, fly, and worms. Identical residues are highlighted in red and similar residues are highlighted in pink. Regions involved in interactions with TMC-1 are depicted as blue bars and the region of ARRD-6 interaction is shown as a yellow bar. Secondary structure elements are shown below the sequence and the location of human CIB2 deafness mutations are indicated above the sequence.

**Extended Data Fig. 9.**
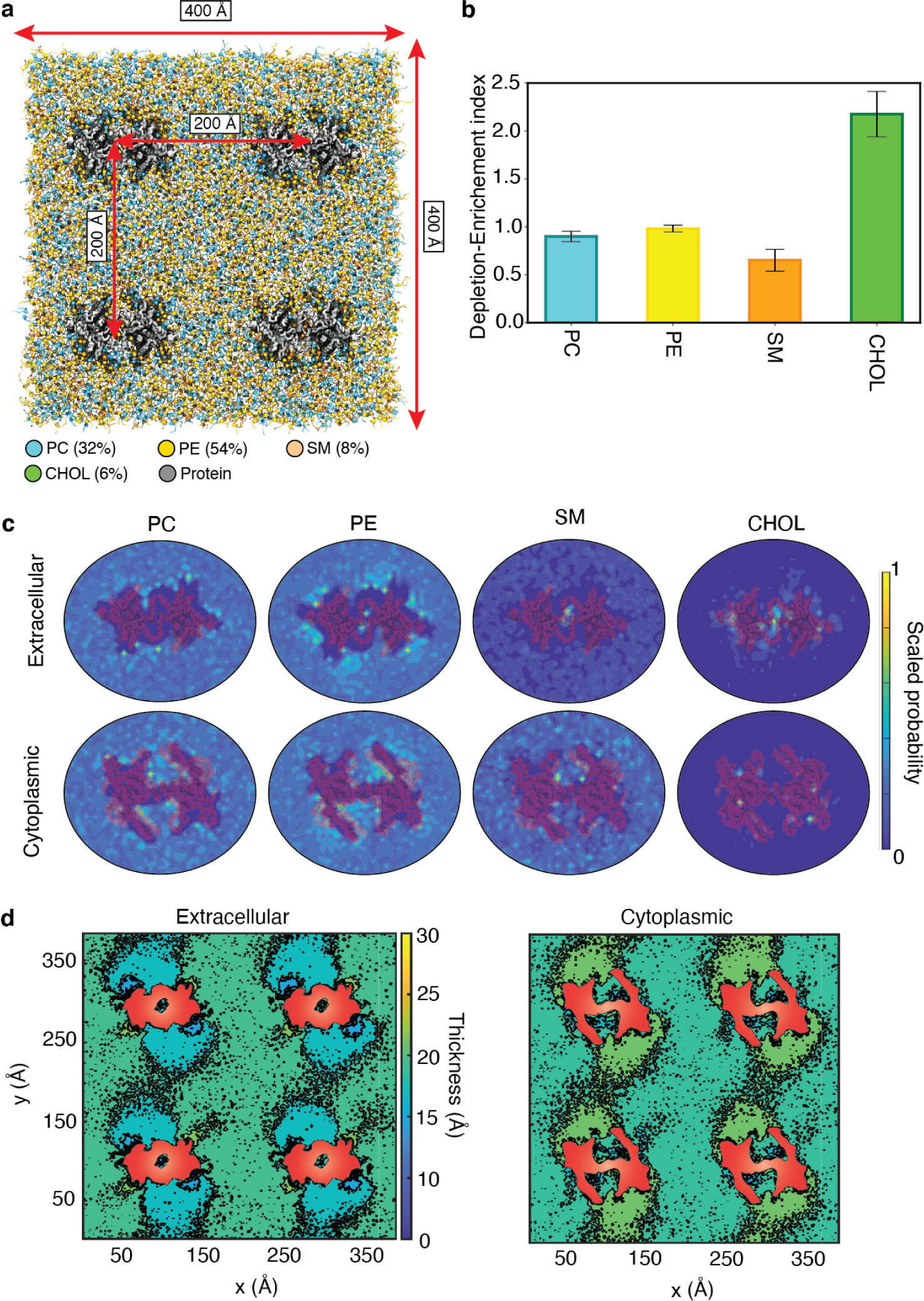
Coarse-grained MD simulations of TMC-1 complex in a membrane bilayer. **a,** Four TMC-1 complexes (gray) in the ‘E’ conformation embedded in a lipid bilayer composed of PC, PE, SM, and cholesterol (CHOL) shown in cyan, yellow, orange, and green, respectively, with a molar ratio of 32:54:8:6. **b,** Enrichment-depletion index of each lipid component in the proximity of the protein. PC and PE densities in the bulk and in proximity of the protein are similar, whereas SM is depleted and CHOL is enriched in the vicinity of the protein relative to their bulk concentrations. **c,** Heatmaps representing distributions of different lipid species around the protein. Each distribution is calculated for the last 4 μs of the trajectory and averaged over all 4 protein replicas. **d,** Lipid bilayer thickness calculated for the extracellular and cytoplasmic leaflets averaged over the last 4 μs of the trajectory. The cross-section of the protein is shown in red. The color scale represents the thickness of each leaflet with blue and yellow corresponding to thinning and thickening, respectively.

**Extended Data Table 1.**
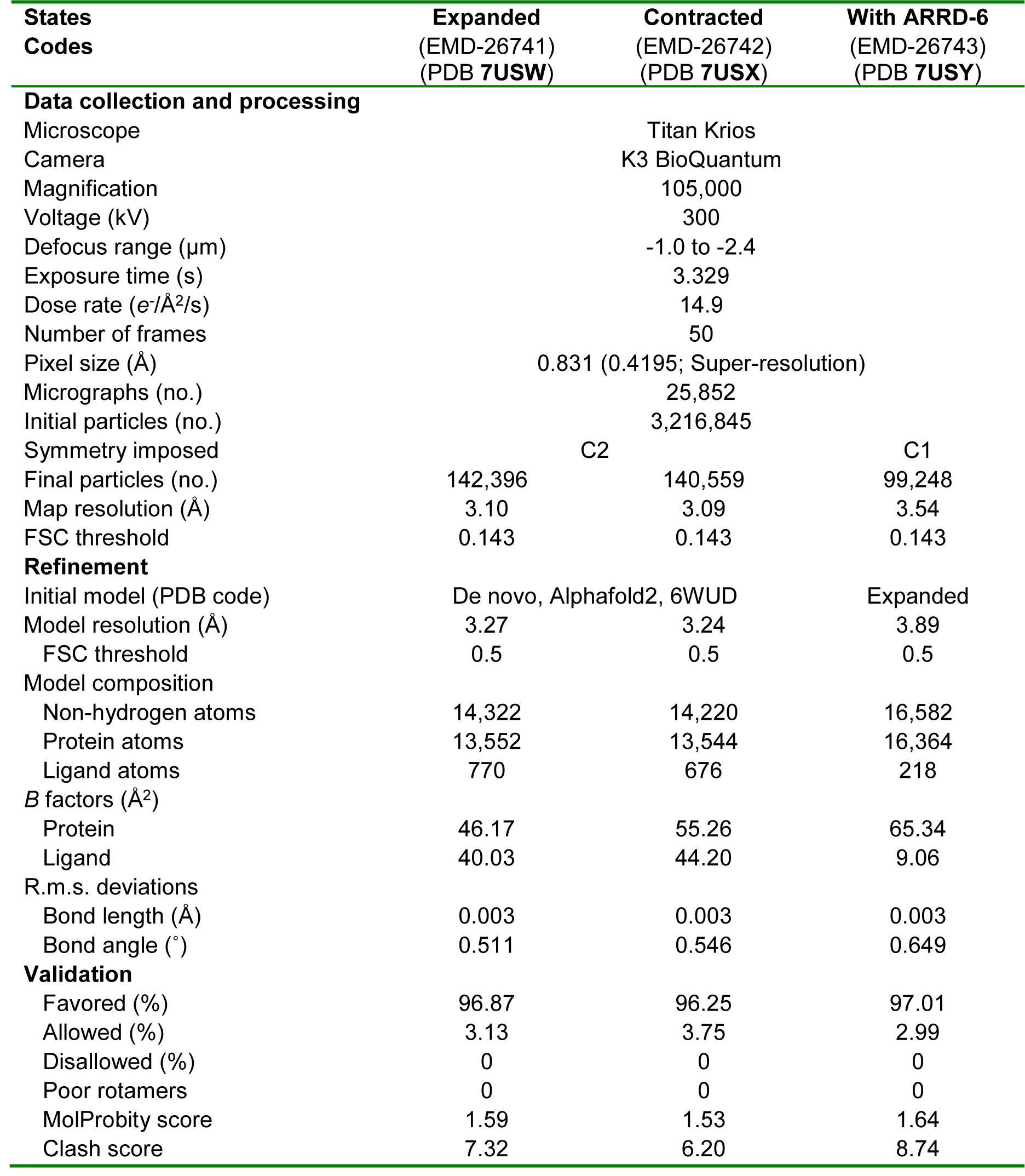
Statistics for 3D reconstruction and model refinement.

**Supplementary Fig. 1.**
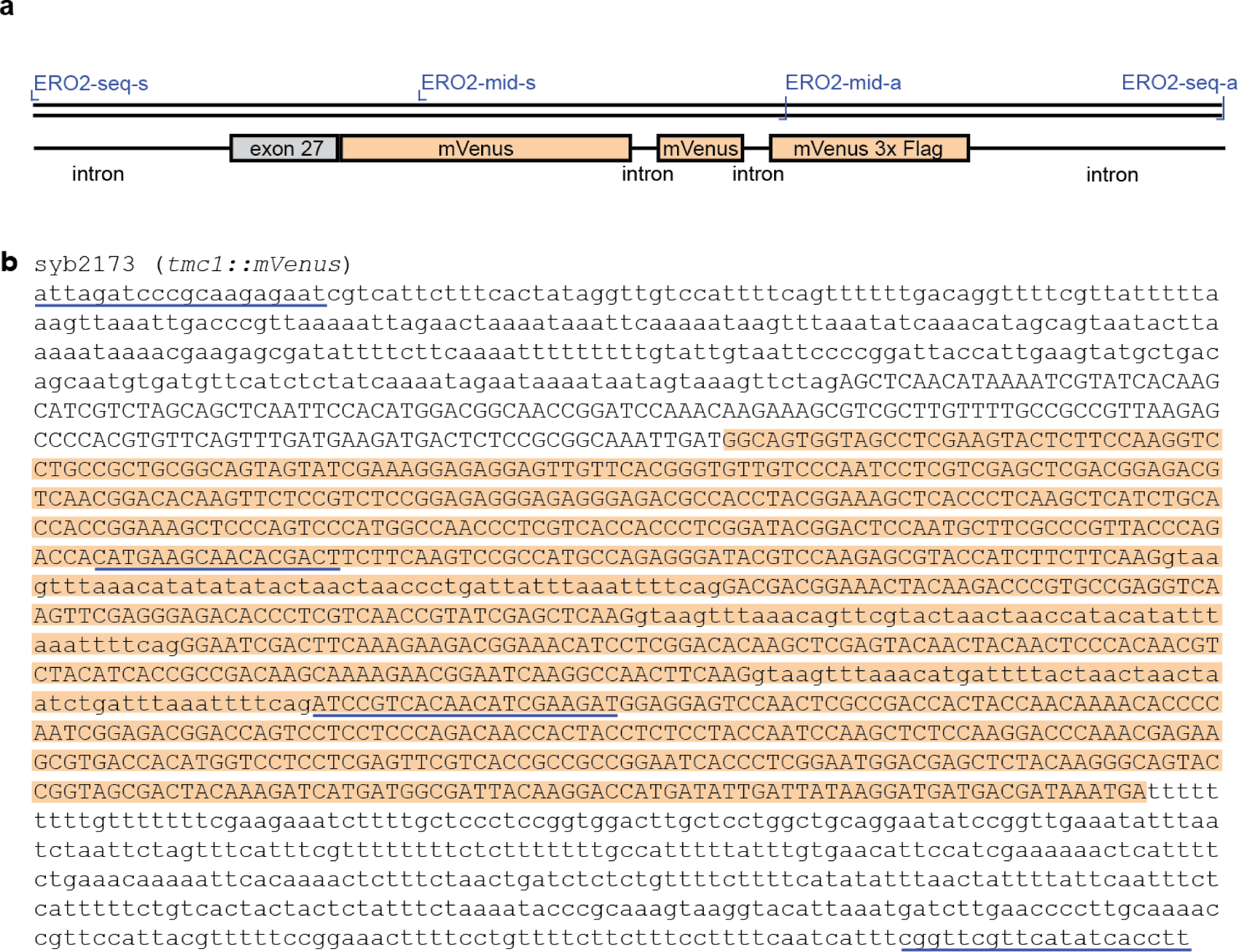
*tmc-1::mVenus* strain. **a**, Schematic diagram of *tmc-1::mVenus* strain. Location of the PCR and sequencing primers relative to the insertion site of 3C precision protease site-mVenus- 3xFLAG prior to the stop codon of *tmc-1*in exon 27. **b,** Sequence of the *tmc-1::mVenus* strain. Introns are in lower case and exons in upper case letters. The 3C precision protease site-mVenus-3xFLAG insert is highlighted in orange. Primer annealing sites for PCR amplification and sequencing of insert are labeled with blue lines.

**Supplementary Fig. 2.**
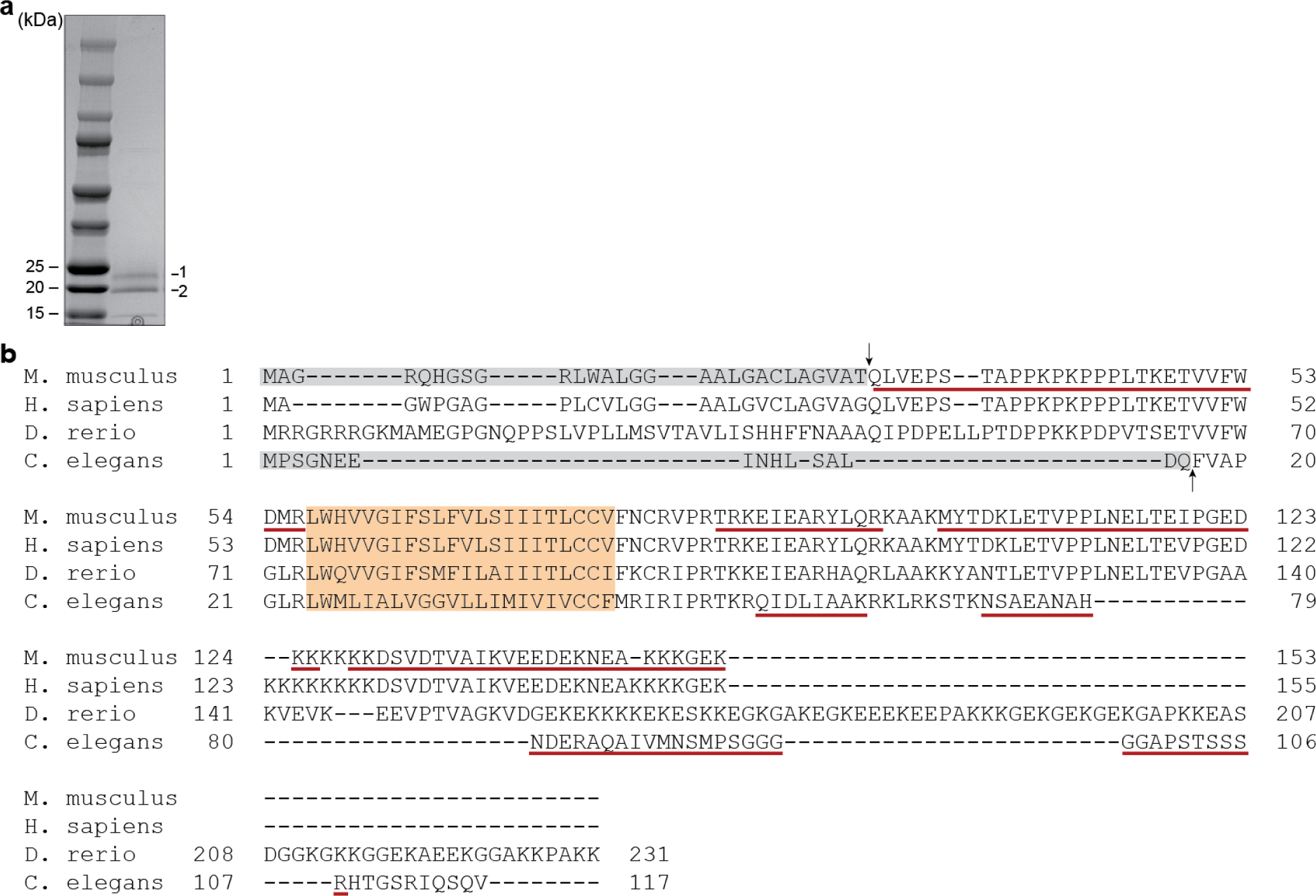
N-terminal sequence of mouse TMIE. **a,** Representative Coomassie staining of sodium dodecyl sulfate-polyacrylamide gel electrophoresis of recombinantly expressed mouse TMIE. Upper and lower bands for mouse TMIE sample submitted for N-terminal sequencing and LC-MS/MS are indicated as 1 and 2, respectively. **b,** Results from N-terminal sequencing of mouse TMIE and mass spectrometry for both mouse and *C. elegans* TMIE. The cleavage site for both mouse TMIE bands was identical and is indicated with an arrow. The identified peptides for both mouse and *C. elegans* TMIE are underlined in red and the signal peptide and transmembrane domain are highlighted in grey and orange, respectively. The peptides that were identified for mouse TMIE were the same in both the upper and lower bands, suggesting the difference in the upper and lower bands is not due to N-terminal or C-terminal cleavage and is most likely due to post translational modification.

**Supplementary Fig. 3.**
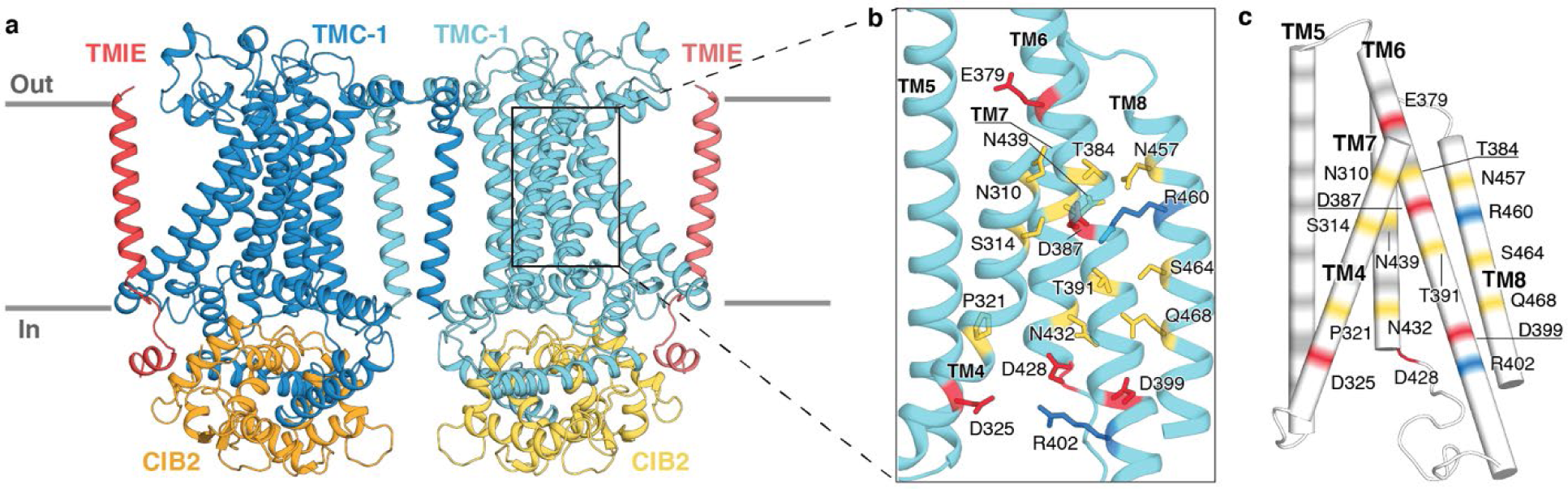
Residue composition of the putative ion conduction pathway of the human TMC-1 complex homology model. **a,** Homology model of human TMC-1 complex: TMC-1 (dark blue and light blue), CIB2 (orange and yellow), and TMIE (red and pink). **b,** An expanded view of the putative ion conduction pathway, highlighting pore-lining residues. Polar (yellow), acidic (red), and basic (blue) residues are shown as sticks. **c**, Electrostatic potential of pore-lining residues are depicted in different colors: grey = nonpolar, yellow = polar, red = acid, blue = basic. Acidic, basic, and polar residues are labeled.

